# Prospects for transcranial temporal interference stimulation in humans: a computational study

**DOI:** 10.1101/602102

**Authors:** Sumientra Rampersad, Biel Roig-Solvas, Mathew Yarossi, Praveen P. Kulkarni, Emiliano Santarnecchi, Alan D. Dorval, Dana H. Brooks

## Abstract

Transcranial alternating current stimulation (tACS) is a noninvasive method used to modulate activity of superficial brain regions. Deeper and more steerable stimulation could potentially be achieved using transcranial temporal interference stimulation (tTIS): two high-frequency alternating fields interact to produce a wave with an envelope frequency in the range thought to modulate neural activity. Promising initial results have been reported for experiments with mice. In this study we aim to better understand the electric fields produced with tTIS and examine its prospects in humans through simulations with murine and human head models. A murine head finite element model was used to simulate previously published experiments of tTIS in mice. With a total current of 0.776 mA, tTIS electric field strengths up to 383 V/m were reached in the modeled mouse brain, affirming experimental results indicating that suprathreshold stimulation is possible in mice. Using a detailed anisotropic human head model, tTIS was simulated with systematically varied electrode configurations and input currents to investigate how these parameters influence the electric fields. An exhaustive search with 88 electrode locations covering the entire head (146M current patterns) was employed to optimize tTIS for target field strength and focality. In all analyses, we investigated maximal effects and effects along the predominant orientation of local neurons. Our results showed that it was possible to steer the peak tTIS field by manipulating the relative strength of the two input fields. Deep brain areas received field strengths similar to conventional tACS, but with less stimulation in superficial areas. Maximum field strengths in the human model were much lower than in the murine model, too low to expect direct stimulation effects. While field strengths from tACS were slightly higher, our results suggest that tTIS is capable of producing more focal fields and allows for better steerability. Finally, we present optimal four-electrode current patterns to maximize tTIS in regions of the pallidum (0.37 V/m), hippocampus (0.24 V/m) and motor cortex (0.57 V/m).

## 1 Introduction

Transcranial current stimulation (tCS) entails sending weak (≤2 mA) currents through the brain via electrodes placed on the scalp (see Reed and Cohen Kadosh, 2018, for a review). Applying direct current (tDCS) has been shown to affect the excitability of stimulated neurons (Nitsche and Paulus, 2000; Dissanayaka et al., 2017), while alternating currents (tACS) can also synchronize neuronal spikes and entrain them to the stimulation frequency (Zaehle et al., 2010; Tavakoli and Yun, 2017). At intensities commonly used for stimulation in humans, tCS does not produce sufficiently high field strengths to directly induce action potentials in neurons (Datta et al., 2009; Rampersad et al., 2014). Mechanisms of tCS action are thought to involve modulation of neuronal membrane potential in a polarity-specific manner with possible effects on synaptic plasticity (Bikson et al., 2004; Jackson et al., 2016). Over the past two decades, tCS has been used extensively in neuroscientific and clinical research, demonstrating both physiological and behavioral effects in fields ranging from working memory (Brunoni and Vanderhasselt, 2014) and visual processing (Plow et al., 2012) to motor recovery after stroke (Lefebvre et al., 2013), and many others (Lefaucheur et al., 2017).

Targets for tCS have classically been superficial cortical regions, and computer simulations with realistic human head models have shown that electric fields induced by tCS are highest in those areas (Datta et al., 2009; Parazzini et al., 2012; Rampersad et al., 2014). Unfortunately, peak tCS fields are rather large and high field strengths may occur far from the target. Use of novel electrodes (Datta et al., 2009; Saturnino et al., 2017) and computational optimization (Dmochowski et al., 2011; Ruffini et al., 2014; Guler et al., 2016) have allowed better targeting and more focal fields, but nonetheless the various forms of tCS still suffer from low focality and low steerability. Deeper brain areas are also desirable targets for electric stimulation, but hitherto these areas have only been targeted with invasive techniques such as deep brain stimulation (DBS) (Follett et al., 2010; Vidailhet et al., 2005). Though recent measurements of intracranial electric fields during in-vivo tACS (Huang et al., 2017) and tDCS (Chhatbar et al., 2018) in humans showed that it may be possible to induce field strengths in deep brain areas sufficient to drive neuromodulation, stimulation of deep areas without concurrent stimulation of the tissue between the electrodes and deep target region is not possible using traditional tCS.

Recently, transcranial temporal interference stimulation (tTIS) was proposed as a method to achieve focal and steerable deep brain stimulation noninvasively (Grossman et al., 2017). Temporal interference has been used in humans since the 1960s for various purposes^1^ including physical therapy (Goats, 1990), electroanesthesia and electronarcosis (Brown, 1975), and has seen considerable recent interest for noninvasive brain stimulation (Grossman, 2018; Melao, 2018). In its basic form tTIS is achieved by applying two high-frequency tACS stimulators to the head with two electrodes connected to each stimulator (Fig. 1a). The two oscillating electric fields interact and result in a single amplitude-modulated field, the envelope of which oscillates at the difference frequency of the two applied fields (Fig. 1b). Temporal interference stimulation is fundamentally based on two major concepts: first, findings indicating that at high frequency, e.g., above 1 kHz, stimulation does not induce a measurable increase in neural firing (Hutcheon and Yarom, 2000; Bikson et al., 2004). If each stimulators’ frequency is in the kilohertz range with the difference between the two frequencies in a physiological range (e.g. *f*_1_ = 2000 Hz, *f*_2_ = 2010 Hz), it could be possible to stimulate neurons with the difference frequency exclusively. Second, the strength of this “interference” field is determined by the *weaker* of the two fields and the alignment between them. Therefore, the peak of the interference field could potentially be deep in the brain, which is not achievable with conventional tCS. Grossman et al. (2017) proposed tTIS as a non-invasive alternative to DBS (i.e., suprathreshold stimulation) and reported tests of tTIS that supported this idea. They performed a series of experiments with anesthetized mice, including in-vivo whole-cell neural recordings, quantification of c-fos expression, and video recordings of motor activity. All experiments confirmed that in mice 1) neurons responded to the difference frequency of two applied fields oscillating at high frequency, and 2) tTIS modulated neural firing in deep areas without doing so in overlying areas. The motor activity experiments additionally showed that 3) different regions of the motor cortex could be selectively stimulated by changing the ratio of input currents, and 4) muscle twitching was achieved at the difference frequency. These results suggest that tTIS can deliver focal, steerable, noninvasive suprathreshold stimulation to both deep and superficial brain areas in mice.

**Figure 1:**
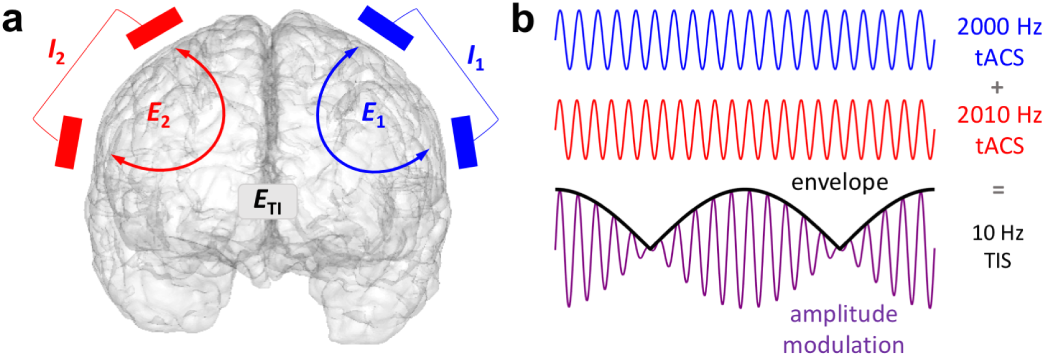
Concept of temporal interference stimulation. **a)** Example arrangement of the two pairs of stimulating electrodes on the scalp, each supplying an oscillating current and producing an oscillating electric field. The intersection of the two fields produces an amplitude-modulated field 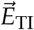. Note that it is not required for the two sets of electrodes to be on opposite sides of the head. **b)** Illustration of two highfrequency oscillations and their sum, which is an amplitude-modulated oscillation with a carrier frequency equal to the average frequency of the inputs and an envelope oscillating at the difference frequency.

While these experiments were promising, effects of tTIS in humans could be vastly different than in mice due to differences in the size, geometry and conductivity of the human head with respect to the mouse. As experiments with human subjects are yet to be published, simulations with a realistic human head model would provide insights into the potential effects of tTIS in humans. A recent study provided the first published simulation results of tTIS in a human head model for two electrode configurations (Huang and Parra, 2019). Here, we present the first: comparison of finite element (FE) simulations of tTIS and tACS in a murine and a human model; simulations of a large set of input current patterns; and investigations of maximal effects along the predominant orientation of local neurons, in addition to effects across all directions. We systematically examined how tTIS-induced electric fields are affected by electrode positioning and input currents to gain an understanding of how these parameters shape the tTIS fields in a human head model.

Finally, we present novel optimization of tTIS current patterns. We investigated to what degree three clinically relevant areas, the left hippocampus, right pallidum, and left motor cortex, can be targeted for tTIS stimulation and how those results compare to tACS. Deep brain stimulation of the pallidum has long been used to treat Parkinsonism (Volkmann et al., 2001; Follett et al., 2010) and dystonia (Coubes et al., 2004; Vidailhet et al., 2005); invasive hippocampal stimulation is a newer therapy to alleviate temporal-lobe epilepsy (Tellez-Zenteno et al., 2006; Velasco et al., 2007) and represents a potential target for Alzheimer’s disease and other forms of dementia; the motor cortex is a commonly used benchmark for novel brain stimulation approaches (Kuo et al., 2013; Fischer et al., 2017). These areas were selected to investigate the potential of tTIS as: 1) a method to noninvasively stimulate deep brain areas, and 2) a more steerable and/or focal alternative for tACS in superficial brain areas. We investigated current patterns and resulting fields for each of these regions optimized over a large number of four-electrode configurations.

Thus this study is a technical exploration of tTIS aimed to better understand its working mechanism and to map the scope of its potential effects. Our results showed that the electric fields induced by tTIS using stimulation currents described by Grossman et al. (2017) are indeed predicted to be high enough to excite neurons in the mouse model, but stimulation currents that are common for human tCS were insufficient in the human model to be able to excite neurons. However, tTIS did achieve field strengths similar to tACS, along with less diffuse, and thus potentially less unwanted, stimulation. Our results suggest that suprathreshold stimulation in humans is not possible with the considered tTIS paradigm but that subthreshold modulation with maxima in deep brain areas is achievable. Thus our results are in direct contrast with both speculations in Grossman et al. (2017), who suggested that tTIS might be an alternative to DBS in humans, and Huang and Parra (2019), who concluded that tTIS does not have any benefits compared to tACS: we conclude that tTIS holds promise as a more steerable and focal version of tACS, making tTIS a potentially valuable noninvasive brain stimulation technique.

## 2 Methods

This section first describes the construction of the murine and human head models employed in this report and then the computational methods that were used to simulate tTIS and tACS. Next, we describe the four studies (one with the murine model and three with the human model) that were performed, and finally the analyses that were carried out on the results of those simulations.

### 2.1 Murine model

An adult male mouse (C57B/L6J) was imaged using a Bruker BioSpec 7.0T/20cm USR horizontal bore magnet (Bruker, Billerica, MA) and a 20G/cm magnetic field gradient insert (ID=12 cm). High resolution T2-weighted images were acquired with a voxel size of 0.117 × 0.117 ×0.250 mm and field of view of 15 × 15× 16.25 mm; this includes the entire skull and cheeks of the mouse, but not the nose. The data set was segmented into skull, cerebrospinal fluid (CSF) and brain using ITK-SNAP v3.4.0 (http://www.itksnap.org); the skin was not included in the model. MR images were aligned and registered to a 3D mouse brain atlas that was segmented and labeled with 20 major anatomical regions using EVA software (Ekam Solutions, Boston, MA). Surface meshes of all compartments were generated using the multiple-material marching cubes (M3C) algorithm (Wu and Sullivan Jr, 2003) with linearization (Sullivan Jr. et al., 2003), which retains the connections between adjoining surfaces. Holes in the mesh were corrected in MeshLab v2016.12 (http://www.meshlab.net), followed by smoothing using the iso2mesh toolbox v1.8 (Fang and Boas, 2009). The surfaces were converted into a linear tetrahedral mesh in TetGen v1.5.1 (Si, 2015) and further refined in the area near the electrodes, resulting in a mesh of length 15.7 mm (anterior to posterior) with 198k nodes, 1.09M elements and a maximum element size of 0.008 mm^3^ (Fig. 2a). The total brain volume was 465 mm^3^. Conductivities were assigned to skull (0.007 S/m), CSF (1.79 S/m) and all brain tissues (0.33 S/m). The left and right motor cortices were identified in the FE model using the aforementioned brain atlas (Fig. 2b).

**Figure 2:**
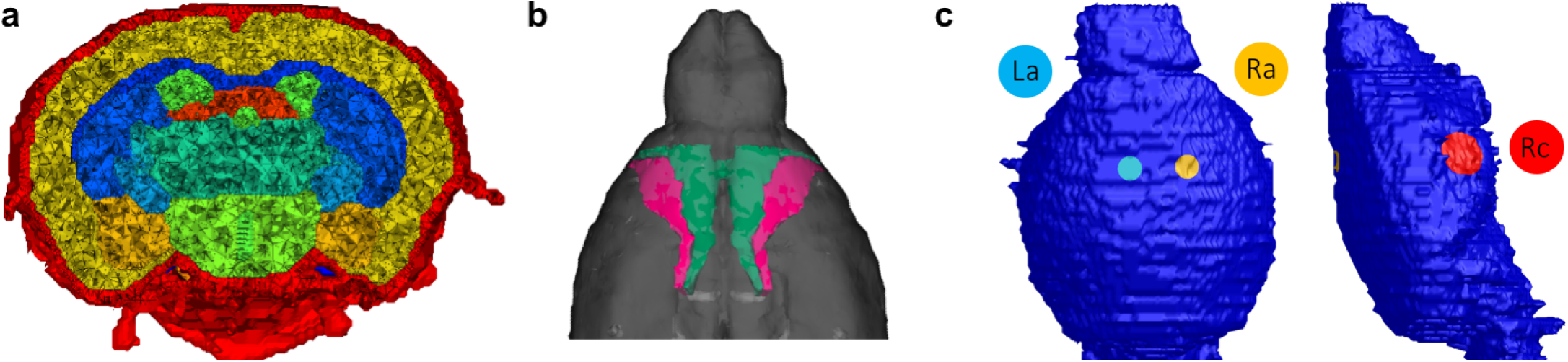
Murine model. **a)** Cut through the finite element model, showing the skull (red), CSF (green), cortex (yellow) and various deeper brain regions. **b)** Location of the left and right primary motor cortex (pink) in the model. **c)** Electrodes on the skull surface of the model, placed based on experiments performed by Grossman et al. (2017) (the left cathode *e*_Lc_ is not visible here; it is placed symmetrically to *e*_Rc_).

### 2.2 Human model

A realistic finite element model was generated from T1-, T2- and diffusion-weighted MR images of a 25-year-old male subject (Fig. 3a). A detailed description of the construction process of an earlier version of the model can be found in Rampersad et al. (2014). All MRI scans were acquired on a 3T scanner (Magnetom Trio, Siemens, Munich, Germany) with a 32-channel head coil, 1 mm^3^ voxel size and a field of view that captured the complete head. The T1-and T2-weighted images were segmented into compartments representing the skin, skull compacta and skull spongiosa using a gray-value based active contour model and thresholding techniques. Eye, muscle and vertebrae segmentations were added manually. The foramen magnum and the two optic canals were modeled as skull openings. The segmentation was then converted into triangular surface meshes and smoothed using CURRY (Compumedics Neuroscan, Charlotte, NC). Segmentation masks of cerebral gray matter (GM), cerebral white matter (WM), cerebellar GM, cerebellar WM, brainstem and ventricles were extracted from a brain parcellation created with Freesurfer (http://surfer.nmr.mgh.harvard.edu). Triangular meshes of the brain surfaces were created and corrected using MATLAB and iso2mesh. Surface meshes of seven skull cavities were created with SimNIBS (Thielscher et al., 2015) using SPM12 (https://www.fil.ion.ucl.ac.uk/spm/) and the CAT12 toolbox (http://www.neuro.uni-jena.de/cat/index.html). All surfaces were then combined into a high-quality 3D Delaunay triangulation via TetGen. This procedure resulted in a mesh consisting of 787k nodes and 4.84M linear tetrahedral elements with a maximum element size of 1.8 mm^3^ in the brain. The total brain volume of the model (excluding cerebellum and brainstem) was 1070 cm^3^. The diffusion-weighted images were processed following previously described methods (Rampersad et al., 2014); conductivity tensors were calculated using the volume-normalized approach (Opitz et al., 2011) and multiplied with the effective conductivity values listed in Table 1 for GM and WM. All other compartments were assigned isotropic conductivities (Table 1)^2^. Segmentation of deep brain structures used as anatomical targets for analysis of our results, the left hippocampus and right pallidum, were produced by Freesurfer and then mapped onto the tetrahedral mesh (Fig. 3b). To construct a target region for the left motor cortex (M1), the location of the cerebral representation of the first dorsal interosseus (FDI) muscle of the right hand was experimentally determined in the volunteer on which the model was based, using single-pulse transcranial magnetic stimulation and electromyography (Rampersad et al., 2014).

**Figure 3:**
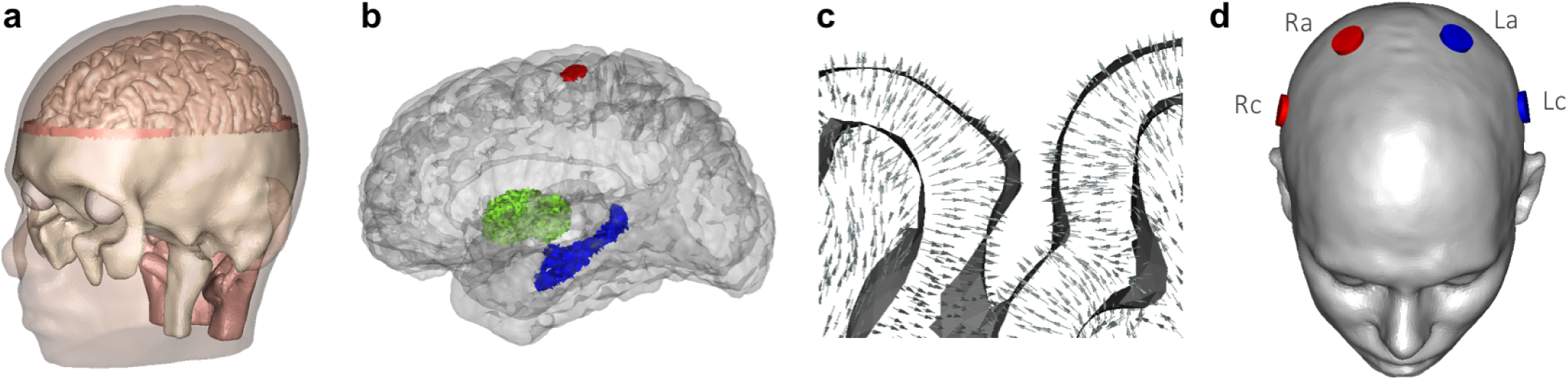
Human head model. **a)** Geometry of the model. **b)** Location of the three selected brain structures: left hippocampus (blue), right pallidum (green) and FDI area of left motor cortex (red). **c)** Section of gray and white matter with arrows representing the preferred direction vectors 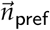. See Fig. S2b for an image of 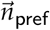 in the whole brain. **d)** Electrodes on the skin surface of the model. Each simulated configuration in Studies 1 and 2 consisted of two electrodes on the left side of the head (blue) that supplied current *I*_1_ and two on the right (red) that supplied *I*_2_. The configuration shown here is the standardized configuration used in Study 1 with electrodes at the C1, C2, C5 and C6 locations of the 10-10 system.

**Table 1:** Tissue compartments in the human head model and conductivities of each compartment. Conductivities denoted with an asterisk were modeled anisotropically based on diffusion-weighted images.

| Tissue | Conductivity (S/m) | Reference |
| --- | --- | --- |
| skin | 0.465 | Wagner et al. (2007) |
| compacta | 0.007 | Akhtari et al. (2002) |
| spongiosa | 0.025 | Akhtari et al. (2002) |
| CSF | 1.65 | Wagner et al. (2007) |
| gray matter | 0.276* | Wagner et al. (2007) |
| white matter | 0.126* | Wagner et al. (2007) |
| brainstem | 0.201* | average GM/WM |
| eye | 1.5 | Nadeem et al. (2003) |
| muscle | 0.4 | Faes et al. (1999) |

### 2.3 Electrode configurations and labeling

We simulated tTIS and tACS with two current sources by placing four electrodes, in a variety of configurations, on the surfaces of the murine and human head models. We can label one current source *I*_1_ and the other *I*_2_, with arbitrary ordering. When connected to a conducting medium through electrodes, *I*_1_ and *I*_2_ produce electric fields *E*_1_ and *E*_2_ in the conductor. The two electrodes supplying *I*_1_ were labeled *e*_1*a*_ (“anode”) and *e*_1c_ (“cathode”), and similarly using *e*_2*a*_ and *e*_2c_ for *I*_2_. Note that since both tTIS and tACS use alternating currents, the anode and cathode labels only refer to the potentials assigned for the initial simulations (as described in Section 2.4). We carried out one study with the murine model and three with the human model, described below. In the mouse study, and Studies 1 and 2 with the human model, two electrodes were placed on the left and two on the right side of the head. For these studies, *I*_1_ and *I*_2_ were denoted by *I*_L_ (left) and *I*_R_ (right) to aid interpretation, and the electrodes were labeled accordingly.

### 2.4 Simulation of *E*_1_ and *E*_2_

We used the Laplace equation, ∇·*σ*∇ Φ = 0, where *σ* and Φ are the electric conductivity and potential respectively, to describe stimulation-induced fields, which implies both quasi-static conditions and linearity (Plonsey and Heppner, 1967). Thus fully simulating stimulation with time-varying current sources only requires solving the equations once with an arbitrary current amplitude, followed by scaling all results to match the stimulation waveform over time. For the results presented here, we were interested in the highest achievable effects and therefore only required the peak values of *E*_1_ and *E*_2_. First, the surface nodes of electrode *e*_1*a*_ were assigned a potential of +Φ_0_ and those of *e*_1c_ -Φ_0_. The Laplace equation was then solved with Dirichlet boundary conditions, Φ = Φ_0_, on the surfaces of *e*_1a_ and *e*_1c_, and Neumann boundary conditions, ∇*σ* Φ·*n* = 0, on the remaining surface, with constant potential across the nodes of each electrode. The system of equations was solved with the FEM solving package SCIRun 4.7 (http://www.scirun.org) using a conjugate gradient solver and Jacobi preconditioner with a maximal residual of 10^*-*10^. From the resulting potential Φ at the nodes of the mesh, the electric field *E* = −∇Φ was calculated in each element and the entire solution was scaled such that the total current flowing between *e*_1*a*_ and *e*_1c_ was equal to the peak amplitude of the desired sinusoidal waveform for *I*_1_. This process was repeated for an input current *I*_2_ between *e*_2*a*_ and *e*_2c_.

### 2.5 Calculation of tTIS and tACS fields from *E*_1_ and *E*_2_

As described in Grossman et al. (2017), at any location 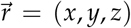, the amplitude of the amplitude-modulated field produced by temporal interference 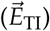 along a direction of interest denoted by unit vector 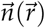 can be calculated from the two component fields 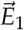 and 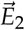 as:

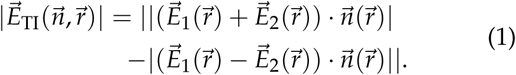

For clarity, we will drop the index 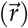 in what follows. From Eq. 1, we can calculate the direction 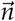 that maximizes the tTIS field strength in a given location, and calculate the size of the field in this direction, which would be the maximal field strength achievable in this location. We will call the local maximum tTIS field strength over all directions 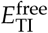 and calculate it as:

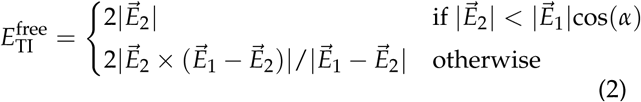

If 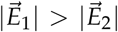, and vice versa otherwise (Grossman et al., 2017). The *α* in this equation denotes the angle between 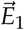 and 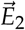, and the equation holds only if *α* is smaller than 90 degrees. Wherever this is not the case, we can flip the direction of one of the fields to ensure this is true. In this case (which is more likely when the electrode pairs are farther apart), the resulting maximum tTIS field strength in the brain, which is what we will present results for below, is a combination of two time points in one oscillation (i.e., some areas reach peak field strength at a different time point than others). The results should be interpreted as the maximum effect over one oscillation. We note, as explained in the introduction, that the size of the temporal interference effect at a given location is determined by both the alignment between the two fields and the amplitude of the smaller field in a complicated interacting fashion.

There is considerable evidence that neurons respond preferentially to stimulation when the field is oriented along the predominant direction of the neuron (or its axon) (Rushton, 1927; Radman et al., 2009a). Therefore, we calculated a preferred direction unit vector 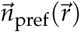 for every brain element in our human model. Specifically, for elements in the white matter, brainstem and cerebellum of the model, the unit vector for element 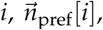, was aligned along the direction of the primary eigenvector of the conductivity tensor calculated from the DTI data in that element. For elements in the gray matter compartment, 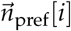 was constructed by interpolating between the nearest normal vectors perpendicular to the GM and WM surfaces in element *i* (Figs. 3c, S1a). Outside of the brain, 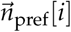 was not defined. The strength of the tTIS field in the preferred direction at each location was then calculated by inserting 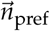 into Eq. 1:

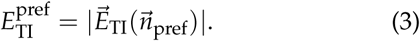

All results for tTIS simulations were compared to the field strengths that would be reached when stimulating with tACS with equivalent parameters. We compared the maximal and preferred tTIS field strengths, 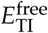 and 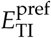, to the corresponding field strengths produced by tACS, 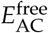 and 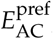, where:

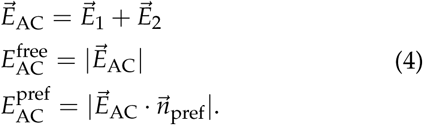

In order to compare tTIS and tACS with identical electrode configurations, two pairs of electrodes stimulating with the same frequency were used here for tACS and the resulting fields were superimposed by a vector sum. Therefore, differently than with conventional tACS, we needed to take polarity into account: flipping one of the vector fields with respect to the other will produce a different result when the fields are summed. As a consequence, in all analyses, Eq. 4 was calculated with 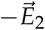 as well, and the configuration that produced the highest field strengths was presented in the results section. Note that for tTIS, polarity does not need to be taken into account, because the two pairs oscillate with different frequencies so their phase difference varies over time regardless of initial electrode assignment; relative polarity determines the phase of the envelope, but does not affect the maximum amplitude.

### 2.6 Steerability

For both the murine and the human model (Studies 1, 2a, 2b), we tested the ability of each stimulation method to steer the peak of the electric field by changing the ratio between input current amplitudes *I*_1_ and *I*_2_. The total current *I*_tot_ was kept constant, while the ratio *R* between *I*_1_ and *I*_2_ was varied:

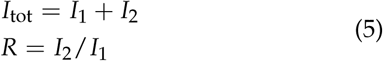

For all simulations, the values used for R were 101 logarithmically spaced values between 0.1 and 10. As we are modeling tCS simulation to be linear with respect to source amplitude, the resulting fields could be scaled linearly with the input currents, so 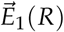 and 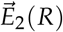 were calculated using 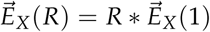.

### 2.7 Simulations with murine model

We followed the specifications of the experiments performed by Grossman et al. (2017) as closely as we could. Four electrodes were embedded into our murine head model (Fig. 2c): two circular electrodes with a radius of 0.5 mm were placed on the skull relative to bregma at AP −1.5 mm and ML −0.5 mm (*e*_La_) or ML +2 mm (*e*_Ra_), which correspond to areas above the forepaw and whisker areas of the left and right motor cortices, respectively. Two electrodes with a radius of 1 mm were placed on the skull near the left (*e*_Lc_) and right (*e*_Rc_) cheeks of the mouse. Electrodes *e*_La_ and *e*_Lc_ formed one current loop, simulating *I*_*L*_ that produced *E*_*L*_, and similarly for *E*_*R*_^3^. Following the experiments reported by Grossman et al. (2017), *I*_tot_ was set to 0.776 mA.

### 2.8 Studies of tTIS in a human head model

Three studies were performed using the human head model described above. These studies first investigate the most basic form of tTIS and then gradually increase the complexity of investigated current patterns and analyses. Study 1 investigates current ratio variations for one configuration; Study 2 examines variations in configurations first for *R* = 1 and later for other ratios. Separating out the effects of these parameters was necessary to fully understand the intricate interplay between tTIS parameters and induced fields. The configurations used in Studies 1 and 2 were based on the experiments by Grossman et al. (2017). By systematically varying the parameters in these models it was possible to study how variations in effects arise. However, these configurations do not necessarily achieve maximal field strengths. In Study 3 we go beyond the standard setup to explore the limits of tTIS-induced field strength and focality.

For all studies, each model was constructed by building four cylindrical electrodes with 1 cm radius, 3 mm height, and conductivity of 1.4 S/m on the skin surface of the head model. For all simulations, *I*_tot_ = 2 mA, a commonly used value for tCS experiments with human subjects (Lefaucheur et al., 2017).

#### Study 1: Standard tTIS and influence of current ratio

To assess the effects of “standard” tTIS with *I*_1_ = *I*_2_, and to investigate the effect of varying the ratio *R* between *I*_1_ and *I*_2_, we first simulated tTIS for one standardized configuration. Based on Grossman et al. (2017), electrodes were placed at the C1, C5, C2 and C6 locations of the 10-10 electrode system (Chatrian et al., 1985), resulting in a configuration with all electrodes in the coronal plane and placed symmetrically with respect to the sagittal plane (Fig. 3d). Electrodes C1 (*e*_La_) and C5 (*e*_Lc_) were used to simulate *I*_L_ and C2 (*e*_Ra_) and C6 (*e*_Rc_) to simulate *I*_R_. Simulations of tTIS and tACS were first performed with *I*_L_ = *I*_R_ = 1 mA. Steerability of the fields was then investigated by performing simulations for a range of current ratios as described earlier.

#### Study 2: Influence of electrode placement

Study 2a investigates several configurations similar to the one used in Study 1 by moving the electrodes horizontally across the head. Electrode locations were chosen from the 10-10 system in order to provide results for configurations that could be easily replicated in experiments. To further investigate the effect of moving electrodes horizontally, a larger set of configurations was created for Study 2b by densely sampling locations on the circumference of the head. While these configurations have less straightforward practical use than the ones used in Study 2a, they allow a more detailed and complete investigation of the relationship between electrode placement and resulting fields. In Studies 2a and 2b, electrode pairs for *I*_1_ and *I*_2_ were moved relative to each other; Study 2c investigates the effect of varying the distance between electrodes *within* each pair by moving electrodes vertically.

2a – We repeated the electrode model construction process described above for four more sets of four electrodes at coronal planes through Fz, FCz, Cz, CPz or Pz (Fig. 7a). More anterior or posterior locations were not considered because of the close proximity of the four electrodes in such cases.

2b – Each configuration consisted of four electrodes on a coronal plane and the set of all configurations was constructed by moving that plane from anterior to posterior in small steps, so that the electrodes were moved around the skin surface (Fig. 8a). To do so, two ellipses were fit to the circumference of the head model parallel to the axial plane, one through Oz and FPz, and one below through C3 and C4. These locations were chosen inferior to those used in Study 2a to cover a larger area of the head than would have been possible with ellipses through the locations used previously. Next, both ellipses were split by the midline into two semi-ellipses. On each of four semi-ellipses, 33 points were placed at angles of [−80 : 5 : 80] degrees with respect to the coronal plane, which after projection onto the skin surface produced 33 possible locations for each of four electrodes: *e*_La_ (top left), *e*_Lc_ (bottom left), *e*_Ra_ (top right) and *e*_Rc_ (bottom right). Finally, 33 electrode configurations were created by selecting the four points corresponding to one angle (one color in Fig. 8a). This resulted in a data set of 3,333 simulated fields (33 configurations times 101 current ratios).

2c – Two top electrodes were placed at CP1 (*e*_La_) and CP2 (*e*_Ra_), close to the standard configuration of Study 1, but behind the ears (Fig. 9a). To create bottom electrodes on both sides of the head, 16 points were placed at vertical distances [15 : 10 : 165] mm below the top electrodes and then projected onto the skin surface. Each pair of bottom electrodes equidistant to the top electrodes was selected as *e*_Lc_ and *e*_Rc_ and combined with *e*_La_ and *e*_Ra_ to form one configuration, resulting in a set of 16 configurations.

#### Study 3: Optimization of four-electrode tTIS

In this study our goal was to find optimal stimulus current patterns to maximize the field strength in three regions of interest (ROIs, described in Section 2.9) while minimizing stimulation outside the ROI, using a fixed set of electrode locations. Direct mathematical optimization of tTIS is a non-convex problem for which it is non-trivial to find unique optimal solutions (see Section 4.6 for more discussion). However, for the case of four electrodes without additional constraints, an exhaustive search method can find globally optimal current patterns in a reasonable amount of time. To maximize the search space, we created a set of 88 electrode positions on our human head model by supplementing 61 electrodes of the 10-10 system with additional rows of electrodes to cover the neck and cheeks (Fig. 10a). A simulation of 1 mA stimulation was performed for each electrode position combined with a “reference” electrode on the bottom of the model and the conductivity of all other electrodes set to 0 S/m. Subtracting any two of these fields provides an approximation for the field produced by the corresponding electrode pair. Using this procedure, 3,828 unique electrode pairs and electric fields were created. Each pair was then combined with every other pair with which it did not have electrodes in common, producing 6,995,670 four-electrode configurations. For each configuration, we calculated tTIS and tACS distributions for 21 current ratios, resulting in a total of 146M input current patterns and simulated electric fields.

### 2.9 Analyses

For each configuration and current ratio simulated in the studies described above, the field strengths 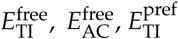 and 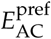 (latter two only in the human model) were calculated in each element of the FE model. From these values, we constructed several parameters of interest. First, in order to remove outliers, the *maximum field strength* in the brain or in a target region was defined as the median value of the top 0.0005% of elements in the volume of interest. To quantify the volume of brain tissue affected by the stimulating field, we calculated the combined volume of all GM and WM elements reaching a field strength larger than a limit value *E*_lim_, which we call the *stimulated volume* or *Vol*_lim_. Since we do not know the exact field strength required to induce effects, results were investigated for a range of *E*_lim_ values, and graphs will be presented for *Vol*_lim_ as a function of current ratio and *E*_lim_. For the human model, we also visualized the spatial distribution of the stimulated volume. For these visualizations, we chose *E*_lim_ = 0.25 V/m based on prior simulation studies. Simulations of tCS with commonly used experimental configurations reached field strengths of 0.15-0.21 V/m in target areas for which experimental studies have shown positive results (Rampersad et al., 2014). To be conservative, we set the lower limit for achieving such subthreshold modulation effects in this report to 0.25 V/m.

In addition to whole-brain analyses, in Studies 1 and 3, stimulation effects were analyzed for three brain structures of clinical or scientific interest. Localization of these structures in the model was described in Section 2.2. In order to compare results for the three structures, regions of interest (ROIs) of equal volume were selected from each structure. For the left motor cortex ROI, a cylindrical region with 1 cm^2^ surface area was selected from the gray matter around the location of the FDI representation in the model; the volume of this area was 129 mm^3^. Spherical areas of equal volume were selected from the head of the left hippocampus and center of the right pallidum.

In Study 3, a large number of configurations was simulated with 21 current ratios. For each *current pattern*, i.e., a combination of electrode configuration and current ratio, and each stimulation modality, we calculated 1) the median field strength (V/m) in the three ROIs, *E*_ROI_, as a measure of effect size, and 2) *Vol*_lim_ (%) for four *E*_lim_ values, as a measure of focality. The goal of this study was to find current patterns that maximize *E*_ROI_ while minimizing *Vol*_lim_. For each *E*_lim_, a Pareto boundary was constructed by first dividing all current patterns into bins of *E*_ROI_ with a width of 0.01 V/m and then selecting the current pattern with the lowest *Vol*_lim_ within each bin. This produced lines of optimal current patterns from which the most appropriate one for a particular experiment can be selected by balancing the requirements of the study in terms of field strength within the ROI and focality. Due to the significant computational cost involved with calculating stimulated volumes for 146M current patterns on a high-resolution mesh, the *Vol*_lim_ values for Study 3 were approximated. The brain compartment of the FE mesh was divided into voxels with edges of 2 mm and the mesh element in the center of each voxel was selected to represent that voxel. The stimulated volume was then calculated by adding the brain volumes within all voxels for which the center element’s field strength surpassed *E*_lim_. Using a smaller electrode set (19 electrodes, 10-20 system) we calculated that the average error introduced by this approach, compared to using all brain elements, was 0.01%.

## 3 Results

In this section we present selected results from all four studies; more detailed results for some aspects of our studies are included in the supplementary material and those figures are denoted by using the letter S before the figure number. In all simulations except those in Study 3, each electrode configuration consisted of one pair of electrodes on the left and one pair on the right side of the head (as in Fig. 1). For these studies, *I*_1_ and *I*_2_ will be denoted by *I*_L_ and *I*_R_ (which produce 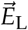 and 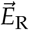, respectively^4^) to aid in the interpretation of results. In the below, “peak(s) of the field” is used to describe area(s) where the field is higher than its surroundings, while “maximum field strength” denotes *the* maximum, i.e., a single value.

### 3.1 Simulations with murine model

This study was an attempt to replicate in silico the in vivo experiments performed by Grossman et al. (2017) for which they reported steerability of tTIS to produce spatially selective induction of activity in the motor cortex of mice, which resulted in twitching of the left or right forepaw or whiskers. We simulated tTIS in a murine FE model designed to mimic their experiments targeting the right motor cortex, with *I*_L_ = *I*_R_ = 0.388 mA. Visualization of the maximum tTIS field strength in any direction 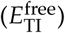 on a plane through the electrodes showed a superficial peak field in the cortex in between the two top electrodes^5^ (Fig. 4a). Stimulation with tACS produced four peaks in the field, one near each electrode (Fig. 4b; the two peaks near the top electrodes appear connected because their values are above the selected limit of the color map), with higher values and larger stimulated volumes than the peaks for tTIS. The maximum 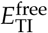 in the brain was 383 V/m and the volume of the brain receiving field strengths of at least 50, 75, 100 or 150 V/m was 19, 7.7, 4.0 and 1.2 mm^3^, respectively (total brain volume: 465 mm^3^).

**Figure 4:**
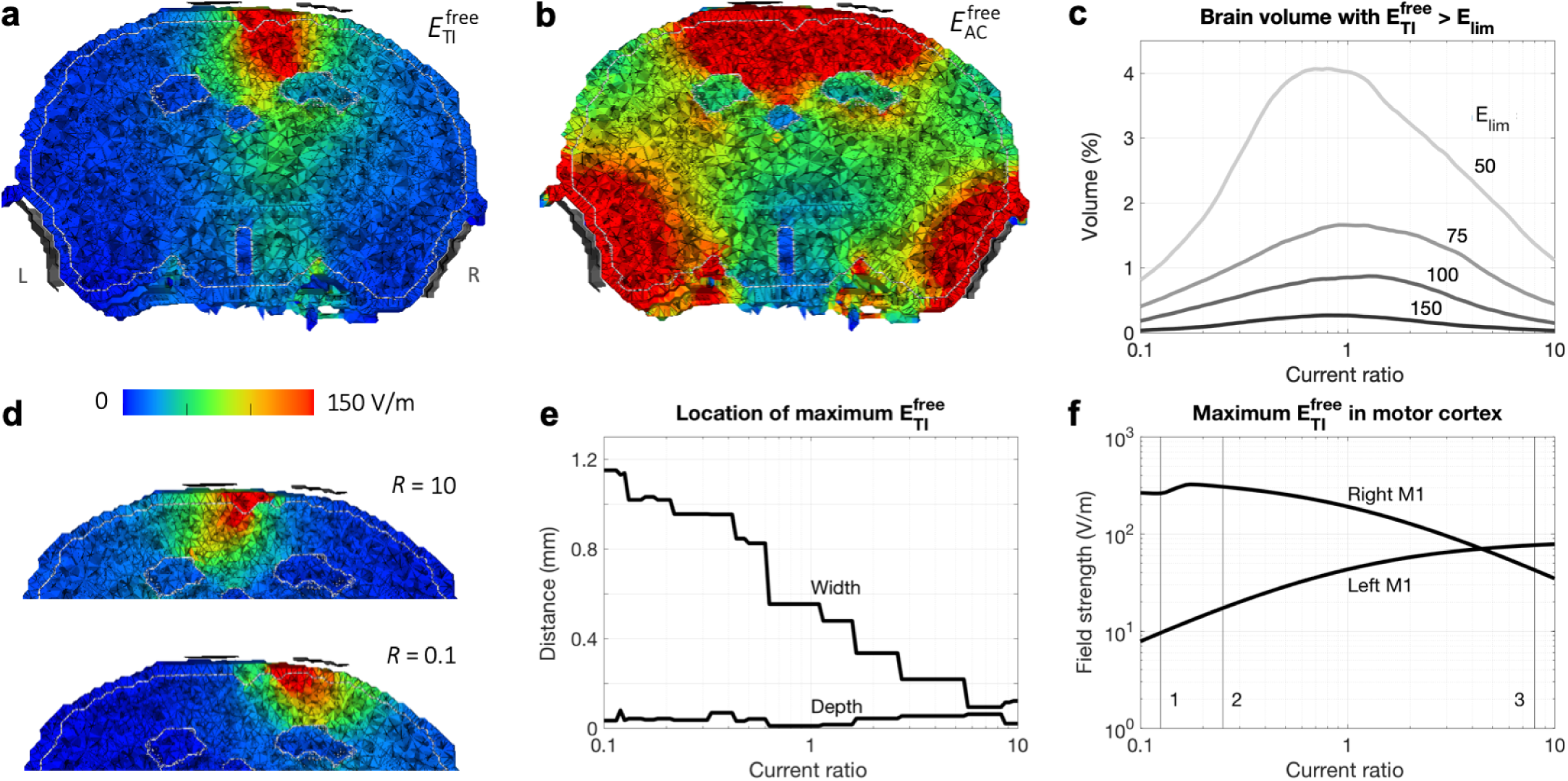
Results of simulations with the murine model. **a**,**b)** *I*_L_ = *I*_R_ = 0.388 mA. Electric field strength on a plane through the electrodes (viewing towards the anterior direction; L and R in panel a indicate the left and right side of the head for all panels) for tTIS (a) and tACS (b). Maxima were 383 V/m for 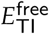 and 38 kV/m for 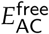. The four electrodes (gray) were displayed at a short distance from the head for visualization purposes. The distinct areas with low field strengths are the ventricles. **c-f)** Current ratios *R* = *I*_R_/*I*_L_ varied from 0.1 to 10 with *I*_L_ + *I*_R_ = 0.776 mA. **c)** Percentage of brain tissue with 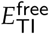 above various limits (limit values indicated in the plot in V/m; total brain volume: 465 mm^3^). **d)** 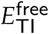 for two extreme values of *R* (compare to *R* = 1 in panel a). See Fig. S2 for corresponding animations with intermediate current ratios for tTIS and tACS. **e)** Distance of the location of maximum 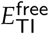 in the brain to the midline (“width”, positive values indicate a location to the right of the midline) and skull surface (“depth”). Lines are not smooth due to the finite size of elements in the model. **f)** Maximum 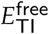 in left and right motor cortex. Numbered lines indicate ratios that elicited the largest movements in the Grossman et al. (2017) experiments for 1: left forepaw; 2: left whiskers; 3: right forepaw and right whiskers.

Increasing the ratio of applied currents (increasing *I*_R_ while decreasing *I*_L_ and keeping *I*_L_ + *I*_R_ at 0.776 mA) moved the peak field towards the left (Fig. 4d, compare to 4a). Since the strength of the tTIS field in any location is determined by the weaker of the two fields in that location, we would expect the tTIS field’s peak to be on the side of the lower amplitude input current. The results thus followed our expectations. When either one of the fields decreases in strength, the tTIS field strength should also decrease. Therefore, stimulated brain volumes decreased as *R* diverged from 1 (Fig. 4c). Since we defined the current ratio *R* as *I*_R_/*I*_L_, the *right* side of each plot (*R* > 1) represents higher currents applied to the *right* side of the head. For each *R*, we calculated the distance between the location of the element with the maximum field strength and either the midline (“width”) or inner skull surface (“depth”). With increasing *R*, the width decreased while the depth stayed fairly constant (Fig. 4e): the peak was effectively moved through the cortex from right to left.

The main stimulation target for the experiments by Grossman et al. (2017) was right motor cortex. For most current ratios, the maximum 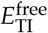 in right M1 was higher than in left M1 (Fig. 4f). For increasing current ratio, maximum 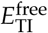 in right M1 first increased and then decreased, peaking at 323 V/m for *R* = 0.17, while maximum 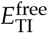 in left M1 increased, surpassing the value in right M1 for *R* > 4.4. The highest 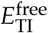 reached in left M1 (78 V/m) was much lower than the highest value for right M1, due to the asymmetric placement of the electrodes closer to right M1, but both values were sufficiently high to expect that stimulation could result in a twitch of the forepaw. The highest field strengths in left and right motor cortex were reached for current ratios that were close to the values reported by Grossman et al. (2017) to elicit the largest movements in right and left forepaw, respectively (numbered lines in Fig. 4f).

### 3.2 Study 1: Standard tTIS and influence of current ratio

Temporal interference stimulation was simulated on a human head model with four electrodes placed in the coronal plane and *I*_L_ = *I*_R_ = 1 mA. Visualizations of the two resulting vector fields *E*_L_ and *E*_R_ (Fig. S1b) show that each field followed a curved path from anode to cathode. Field strength decreased with distance from the center of the electrode pair, with the exception of the corpus callosum area directly above the highly conductive lateral ventricles. The *E*_L_ and *E*_R_ fields combined to produce an interference field following Eq. 2 (Fig. S1c). The 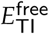 field was strongest at locations where *E*_L_ and *E*_R_ had similar strengths and directions, which generally happened near the centroid of the trapezoid formed by the four electrode locations. Since the temporal interference effect is dominated by the weaker of the two fields at a given location, the direction in which the tTIS field was strongest followed the direction of the field applied to the opposite side of the head. Therefore, in the area below the electrodes, the direction in which tTIS was strongest was nearly perpendicular to the direction in which tACS was strongest (Fig. S1c).

Visualization of 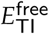 on a plane through the electrodes (Fig. 5c1) shows a distributed peak field near the center of the brain with highest values in the corpus callosum, lower field strengths in superficial regions, and two smaller peak areas in the lower white matter^5^. The maximum 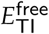 was 0.77 V/m and the volume of the brain receiving field strengths over 0.15 or 0.25 V/m were 181 and 11.7 cm^3^ respectively (total brain volume: 1071 cm^3^). To further investigate the shape of the distribution, we visualized the stimulated area inside the brain (Fig. 6a). The area receiving field strengths over 0.25 V/m consisted mainly of the area above the ventricles, with many smaller disconnected regions.

**Figure 5:**
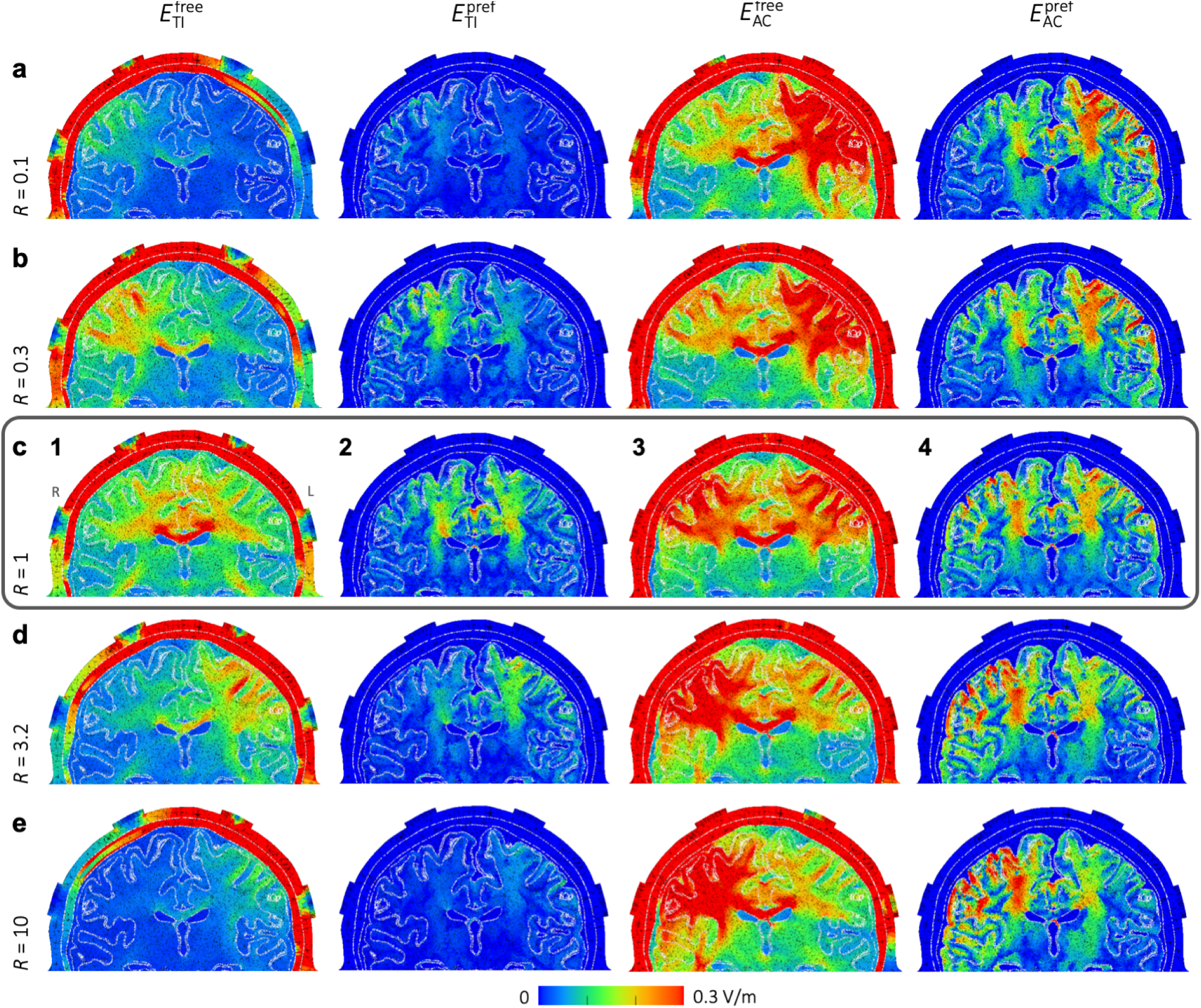
Study 1 – Field strength distributions for tTIS and tACS with various current ratios. Field strengths are displayed on a plane through the electrodes (all placed in the coronal plane), viewing towards the posterior direction (L and R in panel c1 indicate the left and right side of the head for all panels). From top to bottom, the current ratio *R* = *I*_R_/*I*_L_ is increased from 0.1 to 10. Equal current amplitudes, *R* = 1, are shown in the middle row, indicated with a surrounding box. From left to right, 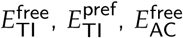 and 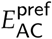 are displayed. Since the preferred direction is only defined for brain elements, all non-brain elements have a value of 0 for plots of 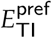 and 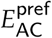. Note that since we are displaying electric field strength, values will be high in areas with low conductivity (such as the skull) and low for highly conductive regions. See Fig. S3 for corresponding animations for intermediate current ratios, and Fig. S1 for visualizations of the directions of *E*_L_, *E*_R_, 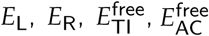 and 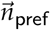.

**Figure 6:**
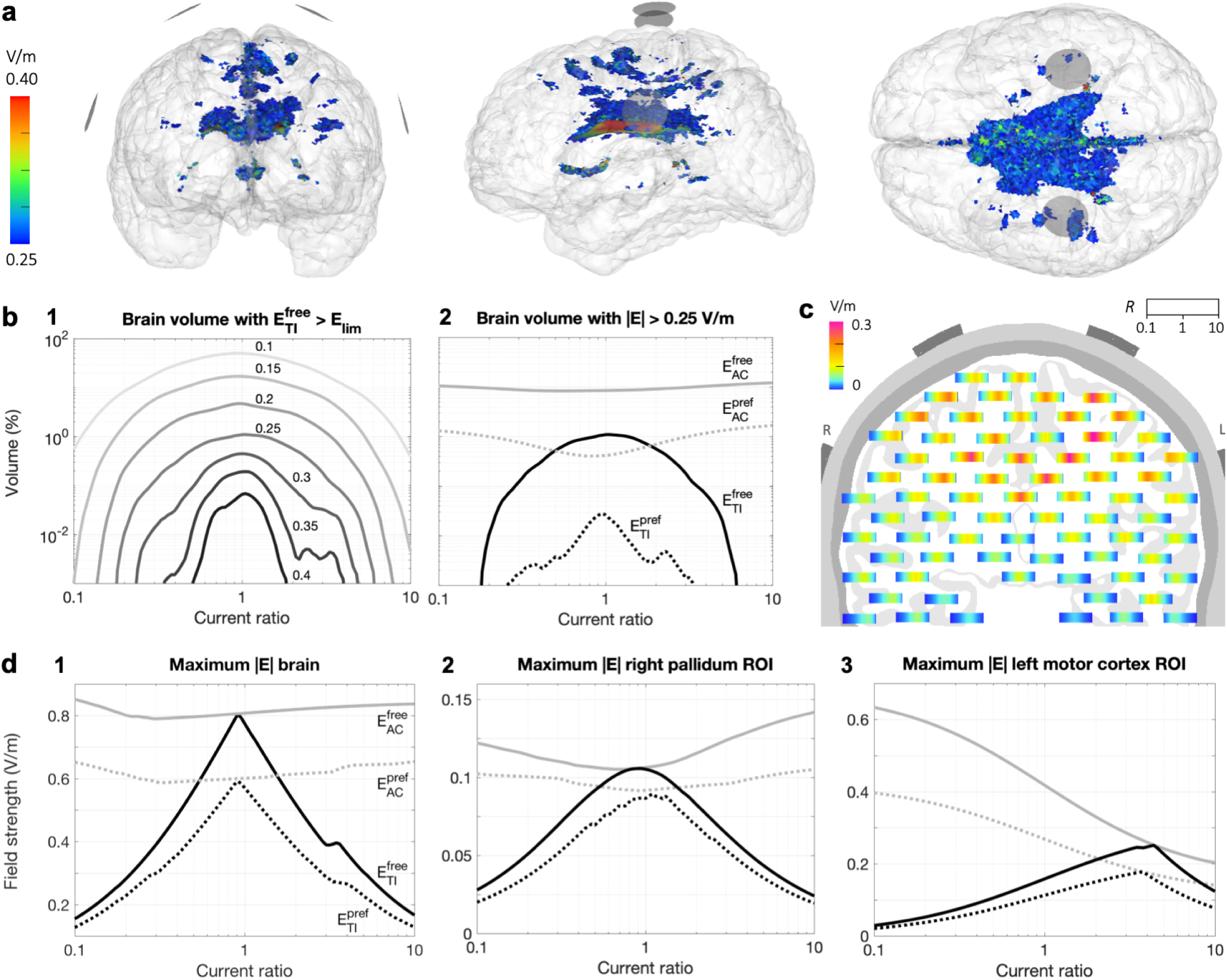
Study 1 – Stimulated brain volumes and maximal field strengths for tTIS and tACS. **a)** Brain volume for which 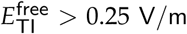 for a simulation with *I*_L_ = *I*_R_ = 1 mA, visualized from the front, left and top, respectively; images are all on the same scale. Electrode surfaces are visualized as gray disks. See Fig. S4 for corresponding animations for other current ratios and other *E*_lim_ values for tTIS and tACS. **b-d)** Results for simulations with current ratios *R* = *I*_R_/*I*_L_ varied from 0.1 to 10 with *I*_L_ + *I*_R_ = 2 mA. **b)** Percentage of brain volume for which 1) 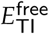 surpasses various limits (values indicated in the plot in V/m), or 2) field strengths surpass 0.25 V/m, for tTIS (black) and tACS (gray) in either a free (continuous lines) or preferred direction (dotted lines). **c)** On a plane through the electrodes, viewing towards the posterior direction (L and R indicate left and right side of the head), each bar displays the local 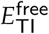 for all current ratios, where horizontal position within each individual bar corresponds to *R* ranging from 0.1 on the left to 10 on the right. **d)** Maximum field strength in the entire brain (1), and in small regions of interest (ROIs) in the right pallidum (2) and left motor cortex (3).

For 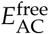 (Fig. 5c3), the field distribution in the center of the brain looked similar to tTIS, with slightly higher field strengths. However, large areas with higher field strengths were located superficially near the electrodes (maximum: 0.81 V/m). It should be noted that with conventional tACS experiments, only two of the four electrodes would be used and therefore there would be only one superficial peak area as opposed to the two seen here.

When only stimulation in the preferred direction (parallel to the neurons) was considered, peak values were lower and peak areas much smaller for both tTIS (Fig. 5c2) and tACS (Fig. 5c4), as would be expected. For 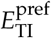, two peak areas were located at the edges of the lateral ventricles and extending upward, and one peak area in the central GM. For this configuration, the tTIS field is strongest near the center of the brain in the vertical direction. Thus, peaks in 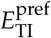 occurred in areas near the center, where the preferred direction is vertical. The preferred direction in the corpus callosum is close to horizontal, leading to low 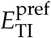 values where 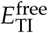 peaked. The same three peak areas can be seen for 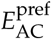, with higher field strengths in the two more distal areas. Additionally, for tACS there were also small peak areas in superficial regions.

Steerability of tTIS was investigated by varying the ratio of input currents *R* = *I*_R_/*I*_L_ from 0.1 to 10 with *I*_L_ + *I*_R_ = 2 mA (Fig. 5, compare the rows of columns 1 and 2). For *R* < 1 (i.e., *I*_R_ < *I*_L_), the peak field moved to the right hemisphere (left side of the image) and for *R* > 1 (i.e., *I*_R_ > *I*_L_), the peak field moved to the left hemisphere, with an additional high-field area in the corpus callosum for a range of *R* values. Similar to the mouse model, the peak was on the side of the head that received the lowest input current, as was expected. For current ratios of 0.17 to 6.6, 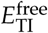 reached values over 0.25 V/m (see Figs. S3 and S4 for results for intermediate *R* values). For tACS, the peak field moved in the opposite direction than for tTIS, since the tACS field is strongest near the highest input current (Fig. 5, columns 3 and 4); field strengths over 0.3 V/m were reached both deep in the brain and in large superficial areas for all *R* values.

Fig. 6 presents several visualizations and plots of stimulated brain volumes and maximum field strengths across all current ratios. For 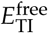, stimulated volumes decreased almost symmetrically as *R* diverged from 1, but volumes were slightly higher for *I*_R_ > *I*_L_ (Fig. 6b1). Field strengths higher than 0.25 V/m or 0.4 V/m were reached for 0.17 < *R* < 6.3 and 0.32 < *R* < 3.6, respectively. For tTIS in the preferred direction, maximum field strengths were approximately 75% of the maximum found in any direction for all ratios (Fig. 6d1), but stimulated volumes were only 1% as large (Fig. 6b2). The effects of current ratio on maxima and stimulated volumes for tACS were minor. A small effect opposite to that for tTIS can be seen; as *R* diverged from 1, the maximum field strength and stimulated volume increased for 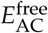 and 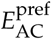. Fig. 6c displays 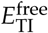 values per current ratio for various locations in the brain. Highest field strengths were reached in an area that follows the curve from the center of each electrode pair to the center of the brain. Close to the midline, highest field strengths were reached for *R* close to 1; towards the left/right side of the brain, peaks were reached for lower/higher current ratios.

For a spherical ROI in the right pallidum, a deep brain structure (Fig. 3b), maxima for 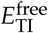 and 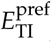 behaved similarly to those in the whole brain, but values were lower and the peak of the curves smoother (Fig. 6d2). The highest 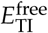 reached in the ROI was 0.11 V/m for *R* = 0.9, i.e., slightly stronger currents supplied to the left side of the head. In a spherical ROI in left hippocampus, the highest 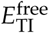 was 0.064 V/m for *R* = 2 (*I*_R_ > *I*_L_). For the FDI area of M1, a superficial structure, effects were highly asymmetric (Fig. 6d3). The highest 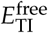 reached was 0.25 V/m for *R* = 4.4, (*I*_R_ ≫ *I*_L_). For 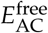, field strengths of 0.14, 0.12 and 0.63 V/m were reached in the pallidum, hippocampus and motor cortex ROIs, respectively, using current ratios at the extrema of the investigated range.

### 3.3 Study 2: Influence of electrode placement

#### Study 2a

For five configurations selected from the 10-10 system (Fig. 7a), simulations were performed with *I*_L_ + *I*_R_ = 2 mA and *R* varied from 0.1 to 10. For equal input currents (*R* = 1), moving the electrodes from front to back lead to a decrease in the maximum 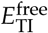 (Fig. 7b) and the stimulated brain volume (Fig. 7c), with the exception of the FC configuration surpassing the F configuration in both measures. These results can be understood by investigating visualizations of the stimulated volumes in the brain (Fig. S5). For all configurations, the stimulated area for tTIS was located deep in the brain, near the midline. While the stimulated area moved from front to back with the electrodes, some of the brain tissue above the ventricles was stimulated in all cases. Because the field is highest in this region, configurations above it (F, FC, C) had the highest peak values and largest stimulated volumes, with a configuration centered above the ventricles (FC) producing the highest values.

**Figure 7:**
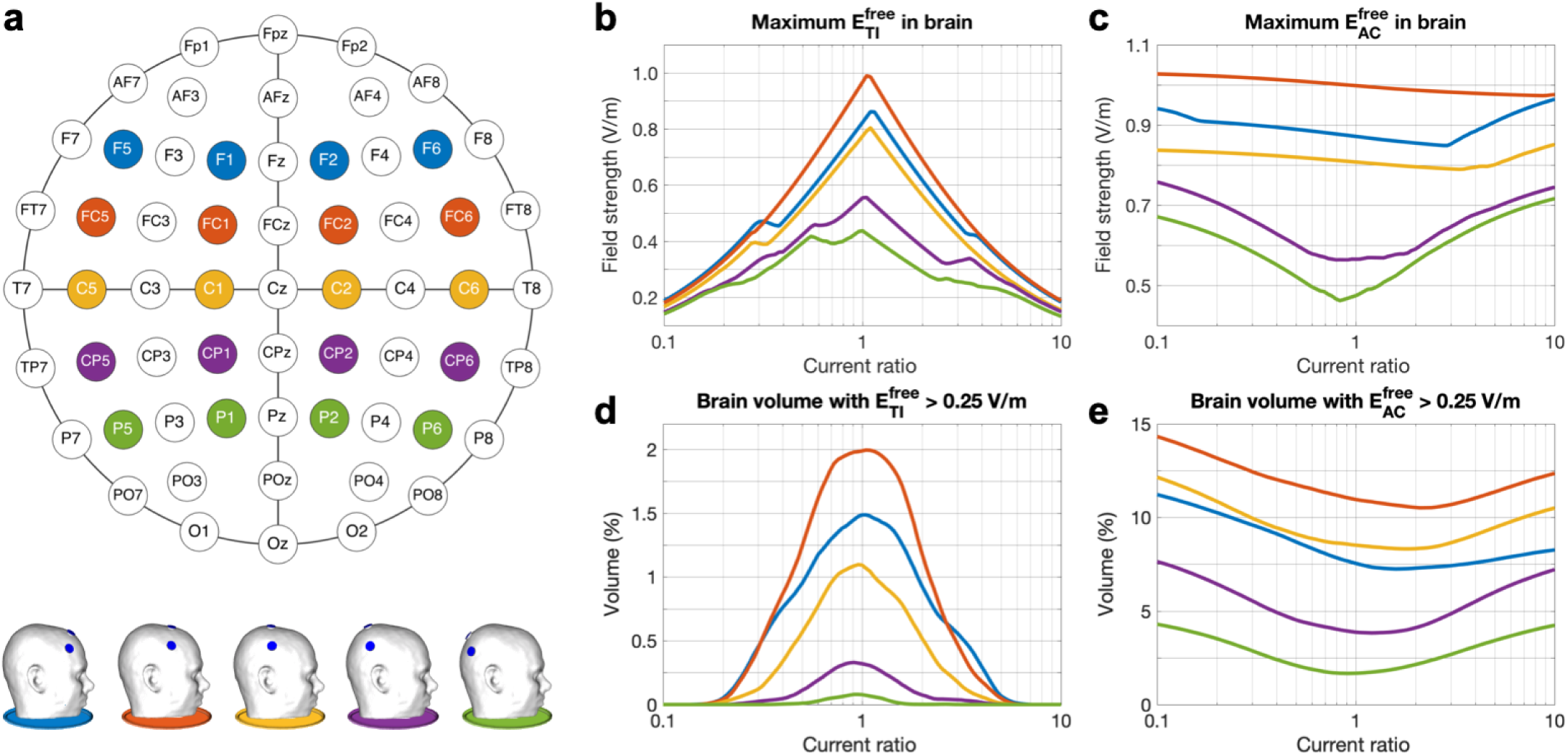
Study 2a – Simulations of tTIS and tACS with five standardized electrode configurations and various current ratios. **a)** Schematic of the 10-10 system with the five configurations marked in different colors (top) and side view of the five head models (bottom). **b**,**c)** Maximum field strength in the brain, and **d**,**e)** percentage of brain volume stimulated at > 0.25 V/m for 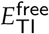 (b,d) and 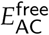 (c,e). Line colors match the schematic and models shown in panel a. See Fig. S5 for animations of stimulated brain volumes (similar to Fig. 6a) for tTIS and tACS for all configurations and current ratios.

The effect of varying the current ratio was consistent across all configurations: the maximum 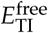 in the brain (Fig. 7b) and the stimulated brain volume (Fig. 7c) decreased strongly as *R* diverged from 1, while an opposite but much smaller effect occurred for 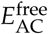 (Fig. 7d,e). Small differences in the shape of the curves between configurations and between the left and right hemisphere are mostly due to differences in cortical folding across the brain. Spatially, the stimulated volumes for tTIS moved off-center for *R* diverging from 1 (Fig. S6). For 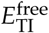 in the most frontal configuration, superficial GM areas were also reached with fields over 0.25 V/m. For tACS, the stimulated volumes were two large superficial regions beneath the electrodes and a smaller deep area above the ventricles. When all configurations and current ratios are combined, tTIS with 2 mA total current was able to reach 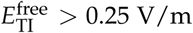 V/m in 13% of the WM volume and 1.7% of the GM volume; volumes for 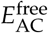 were 34% of WM and 54% of GM.

#### Study 2b

Simulations with 33 parallel electrode configurations were performed with *I*_L_ + *I*_R_ = 2 mA and *R* varied from 0.1 to 10 (Fig. 8a). Maximum 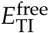 values in the brain and stimulated volumes were highest for *R* close to 1 (Fig. 8b,d), and both quantities increased for more frontal configurations, in agreement with the results for Study 2a. Here, we see that this effect is continuous and smooth across a wide range of angles and persists until the electrodes almost touch in the front. Towards the back, however, the field strength and stimulated volume showed a slight increase again when the electrodes got closer. Visualizations of the stimulated brain area (Fig. S7) demonstrate that higher field strengths and larger volumes were reached when the electrodes were placed on narrower parts of the head. In these locations, the peaks of *E*_R_ and *E*_L_ were close together, so that temporal interference (determined by the weakest of the two fields) was larger, while electrodes placed on wider parts of the head were further away from the point of maximal interference (the midline), so each field, and the corresponding tTIS field produced, was smaller. The anterior peaks were much larger than the posterior peaks, likely due to the difference in curvature of the head (see Fig. S7, side view) and the relative distance to the lateral ventricles. For 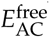, field strengths and stimulated volumes also increased towards the front, but similar values were reached across all current ratios (Fig. 8c,e).

**Figure 8:**
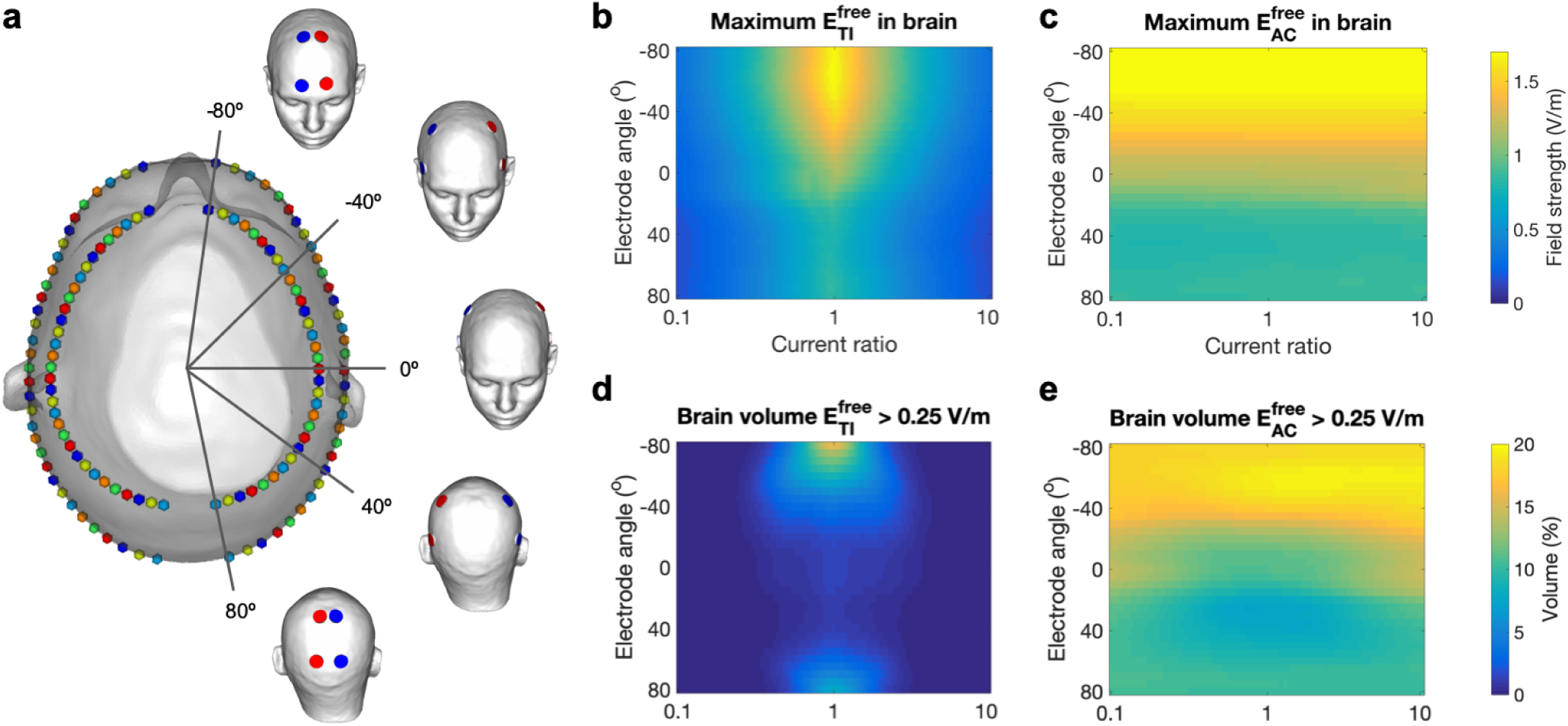
Study 2b – Simulations of tTIS and tACS with a densely sampled set of parallel electrode configurations and various current ratios. **a)** Visualization of the design of the 33 configurations on the head model with five configurations highlighted (red: *I*_*L*_; blue: *I*_*R*_). On a top view of the head model, each configuration is represented by four identically colored spheres. The four electrodes of one configuration were placed symmetrically around the midline with equal angles to the coronal plane; the set of four was moved from anterior to posterior by varying the angle between the electrode locations and the coronal plane from −80 to 80 degrees. **b**,**c)** Maximum field strength in the brain, and **d**,**e)** percentage of brain volume stimulated above 0.25 V/m for 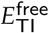 (b,d) and 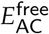 (c,e). Results are displayed on a 2D plot with ratio *R* on the horizontal axis and coronal plane angle in degrees on the vertical axis. See Fig. S7 for animations of stimulated brain volumes (as in Fig. 6a) for tTIS and tACS for all configurations.

#### Study 2c

Effects of moving the electrodes vertically were investigated through simulations with 16 electrode configurations with all electrodes in a vertical plane through CP1 and CP2, with *I*_L_ = *I*_R_ = 1 mA (Fig. 9a). For both tTIS and tACS, for free and preferred directions, the maximum field strength in the brain and the stimulated brain volume increased with electrode distance. As the two electrodes of one electrode pair move closer together, more current is shunted through the skin and skull and less current enters the brain. This results in lower electric fields throughout the brain. Note that while the opposite seemed to happen in Study 2b (field strengths increased when electrodes were moved towards each other), in that study the two pairs of electrodes (oscillating at different frequencies) were moved closer together, which increased the interaction that created the tTIS field. By contrast, in Study 2c, the two electrodes in one pair (oscillating at the same frequency) were moved closer together. As the bottom electrodes were moved downwards, the distribution of the 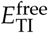 field changed only slightly (Fig. S8); the main effect of moving the electrodes was that the entire field was scaled up. When the electrodes passed below the temporal lobe, the field strength remained constant in the cerebrum and increased in the cerebellum. While maximum field strengths for tTIS and tACS were almost equal for all distances, stimulated volumes for tACS were much larger, suggesting that tTIS produced a more focal field.

**Figure 9:**
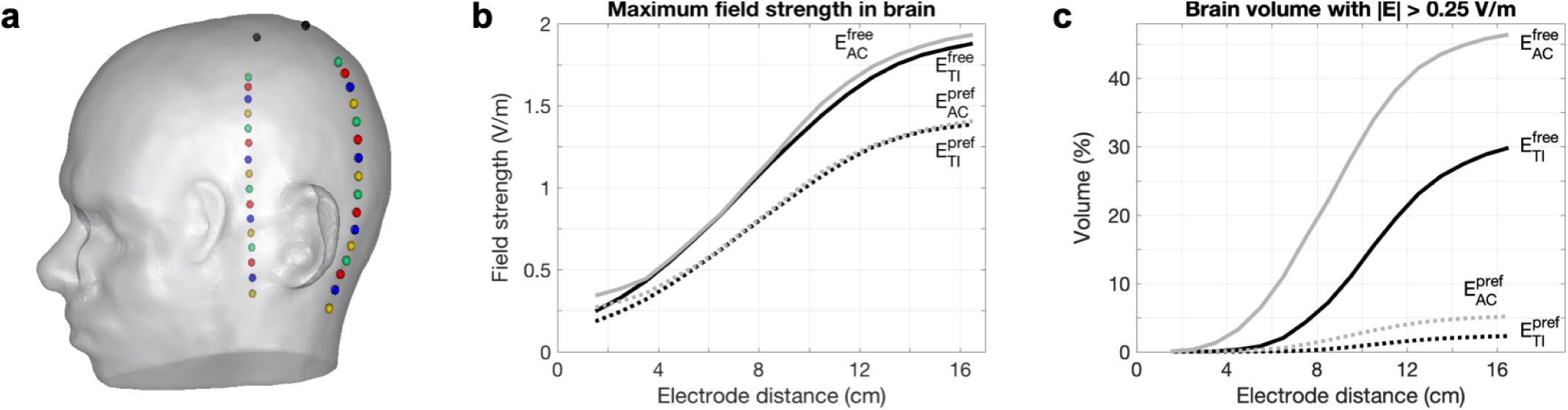
Study 2c – Simulations of tTIS and tACS with electrode locations varied vertically across the head surface. **a)** Visualization of the design of the 16 configurations on the head model. Each configuration consisted of the same two top electrodes (black spheres) and one pair of bottom electrodes (identically colored spheres) placed underneath. **b)** Maximum field strength in the brain, and **c)** percentage of brain volume with field strengths above 0.25 V/m, for tTIS (black) and tACS (gray) in either a free (continuous lines) or preferred (dotted lines) direction. See Fig. S8 for animations of tTIS field strength distributions (as in Fig. 5) and stimulated volumes in the brain (as in Fig. 6a) for all configurations.

### 3.4 Study 3: Optimization of four-electrode tTIS

Simulations of tTIS and tACS were performed with nearly 7M four-electrode configurations and 21 *R* values from 0.1 to 10 with *I*_tot_ = 2 mA. Pareto boundaries of optimal target field strength (*E*_ROI_) and minimal stimulated volume (*Vol*_lim_) were constructed for small ROIs in the head of the left hippocampus, center of the right pallidum, and FDI area of the left motor cortex. Results for four values of *E*_lim_ for all ROIs can be found in Fig. S9; results for *Vol*_0.25_ for the right pallidum are shown in Fig. 10. These Pareto lines were constructed by grouping all current patterns into bins of similar *E*_ROI_ and finding the lowest *Vol*_lim_ within each bin. All results discussed below refer to this set of optimal (i.e. minimal *Vol*_lim_ for a certain *E*_ROI_) current patterns.

**Figure 10:**
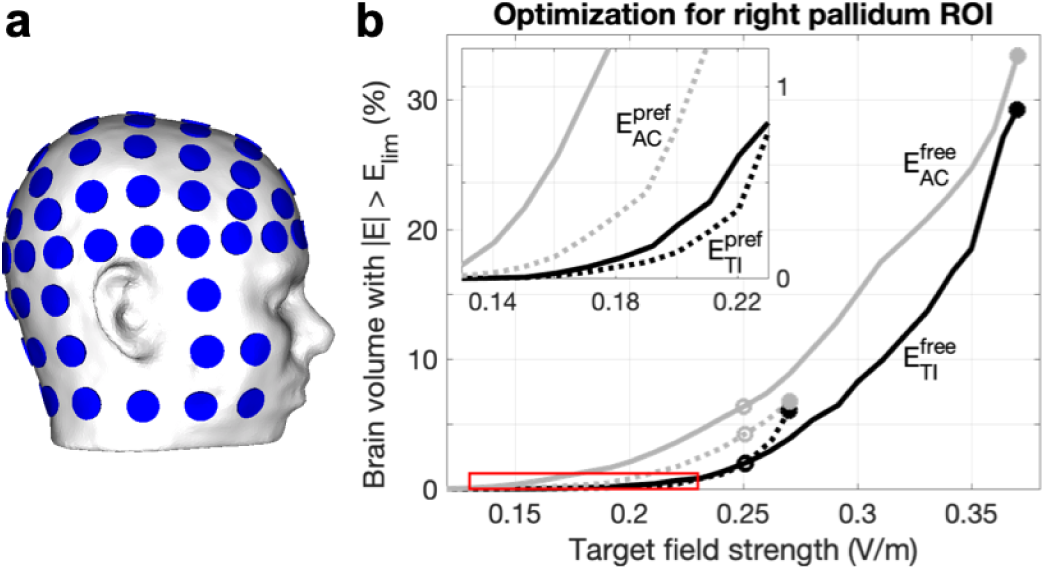
Study 3 – Pareto boundaries of optimal field strength and focality for four-electrode tTIS and tACS in the right pallidum. **a)** A set of 88 electrodes was used to perform an exhaustive search optimization over 146M current patterns. **b)** From this set, these lines present the minimum unwanted stimulation (volume of brain tissue stimulated over 0.25 V/m, *Vol*_0.25_) achievable as a function of median field strength in the target region of interest (ROI, *E*_ROI_), for tTIS (black) and tACS (gray) in either a free (continuous lines) or preferred (dotted lines) direction. From these lines, the most suitable current pattern can be selected to achieve a specific experimental goal. Circles indicate current patterns that minimized *Vol* _0.25_ while either maximizing *E*_ROI_ (filled) or reaching at least 0.25 V/m in the ROI (open); more detailed results for these current patterns can be found in Table 2 and Fig. 11. The red box indicates the location of the inset.

Stimulated volumes increased with *E*_ROI_ for all ROIs, stimulation types, and *E*_lim_ values (Fig. S9). Comparing tTIS to tACS, *Vol*_lim_ was smaller for tTIS for all *E*_lim_ values for all *E*_ROI_ for the two deep ROIs. For motor cortex, *Vol*_lim_ was generally higher for tTIS than for tACS for all *E*_ROI_ for stimulation in any (free) direction, but was lower for *E*_ROI_ < 0.20 V/m for stimulation in the preferred direction. Comparing free to preferred direction, *Vol*_0.25_ was always larger for *E*^pref^ than for *E*^free^ for the hippocampus and M1 ROIs, for both tTIS and tACS. For pallidum, however, *Vol*_0.25_ was smaller for *E*^pref^ for most values of *E*_ROI_.

For each ROI and stimulation type, we selected two current patterns from the Pareto lines with *E*_lim_ = 0.25 V/m to investigate further: 1) the pattern that maximized *E*_ROI_, and 2) the pattern that minimized *Vol*_0.25_ for *E*_ROI_ > 0.25 V/m. First, the optimal results for the highest *E*_ROI_ bin (filled circles in Figs. 10, S9) were investigated to explore how field strength in an ROI can be maximized with tTIS and how this differs from tACS. Maximal *E*_ROI_ values were 0.24–0.56 V/m for 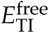 and 0.17– 0.26 V/m for 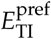 (Table 2). Maximal 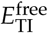 for M1 was 1.5–2.3 times higher than for the two deep targets, but 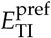 values for M1 and pallidum were similar. Higher field strengths were reached in the pallidum compared to the hippocampus, which could be due to the pallidum ROI being located anterior and superior to the hippocampus ROI. Maximal *E*_ROI_ for 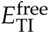 was 2.9–5.4 times larger than the values reached with the configuration used in Study 1 that was based on Grossman et al. (2017). Comparing tTIS to tACS, maximal *E*_ROI_ were nearly equal for the two deep targets, but were 1.1 (*E*^free^) and 1.2 (*E*^pref^) times higher for tACS in M1. For these optimal current patterns, i.e. the ones producing the maximal *E*_ROI_ on the Pareto boundary, stimulated volumes (*Vol*_0.25_) were 15– 46% for *E*^free^ and 2–7% for *E*^pref^ for tTIS and tACS.

**Table 2:** Study 3 – Pareto-optimal results for tTIS and tACS of three target regions. From a large set of current patterns, Pareto boundaries were constructed that maximize field strength in a small target region (*E*_ROI_) while minimizing the stimulated brain volume (*Vol* _0.25_) (Fig. 10). The table lists the maximum achievable *E*_ROI_ on this boundary (filled circles in Fig. 10). Percentages listed indicate the percentage of stimulated brain volume for the current pattern that minimizes *Vol* _0.25_ with *E*_ROI_ *>* 0.25 V/m (open circles in Fig. 10). For the hippocampus target not all stimulation types reached this value, so *E*_ROI_ *>* 0.17 was used. The value of 0.00% in the table indicates a percentage lower than 0.005.

| Target | $E_{TI}^{free}$ | $E_{AC}^{free}$ | $E_{TI}^{pref}$ | $E_{AC}^{pref}$ |
| --- | --- | --- | --- | --- |
| **Left hippocampus** |  |  |  |  |
| Maximum $E_{ROI}$ (V/m) | 0.24 | 0.25 | 0.17 | 0.18 |
| $Vol_{0.25}$ GM (%) | 0.00 | 0.05 | 0.70 | 1.62 |
| $Vol_{0.25}$ WM (%) | 0.69 | 2.81 | 2.67 | 5.09 |
| **Right pallidum** |  |  |  |  |
| Maximum $E_{ROI}$ (V/m) | 0.37 | 0.37 | 0.27 | 0.27 |
| $Vol_{0.25}$ GM (%) | 0.09 | 2.16 | 0.67 | 2.06 |
| $Vol_{0.25}$ WM (%) | 3.64 | 9.89 | 3.02 | 6.11 |
| **Left motor cortex** |  |  |  |  |
| Maximum $E_{ROI}$ (V/m) | 0.56 | 0.60 | 0.26 | 0.31 |
| $Vol_{0.25}$ GM (%) | 0.05 | 0.05 | 0.57 | 0.60 |
| $Vol_{0.25}$ WM (%) | 0.18 | 0.10 | 1.96 | 0.96 |

Next, we investigate what the above-mentioned optimal current patterns look like (Fig. S10). For all three ROIs, the current pattern for 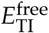 consisted of two nearparallel electrode pairs with minimal distance between the pairs and *R* = 1. The two pairs being parallel maximized the directional agreement between the two fields, while the pairs being close to each other maximized the minimum field strength of the two pairs (as noted, the tTIS field is determined by the lowest field strength of the two fields and the angle between the fields). Placement of electrodes within a pair on opposite sides of the ROI maximized the target field strength created by one pair; for the two deep targets, the electrodes within each pair were far apart so that sufficient current could reach the ROI; for the motor cortex, the highest *E*_ROI_ was achieved with both electrodes close to the target. This resulted in higher field strengths for superficial as compared to deep ROIs. The optimal current patterns for tACS were similar to those for tTIS, but with *R* ≠ 1. We note that, if focality were ignored, maximal *E*_ROI_ for tACS would be reached with similar electrode pairs as found here, but with *R* = 10 (or 0.1), because this would make a configuration of two electrode pairs almost equivalent to a single bipolar configuration. Given a limited amount of current without additional constraints, theoretically a bipolar configuration would always reach maximal field strength for tACS.

The current patterns described above would be optimal for an experiment in which the goal were to stimulate the target as strongly as possible, with minimal *Vol*_0.25_ at that field strength. For an experiment in which focality is more important, one could instead choose a current pattern that achieves at least an effective field strength in the target and produces a much lower *Vol*_0.25_. For this purpose, we selected the current patterns that produced the lowest *Vol*_0.25_ for *E*_ROI_ > 0.25 V/m (open circles in Fig. 10, S9). For the hippocampus ROI not all stimulation types reached this value. Therefore we used *E*_ROI_ > 0.17 V/m for this target to still be able to compare focality across stimulation types. For this second set of optimal current patterns, *Vol*_0.25_ values were calculated for GM and WM separately (Table 2). In all cases, the percentage of stimulated WM was larger than the percentage of stimulated GM. Stimulated volumes for M1 were generally smaller than for the deep targets, for both tTIS and tACS. Comparing tTIS to tACS, the stimulated volumes for tACS were always larger for hippocampus and pallidum: *Vol*_0.25_ was 10–23 (GM) and 2.7–4.1 (WM) times larger for 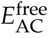, and 2.3–3.1 (GM) and 1.9–2.0 (WM) times larger for 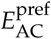. In contrast, for motor cortex *Vol*_0.25_ values were similar in GM and larger for tTIS in WM, though differences between stimulation types were smaller than for the deep targets: *Vol*_0.25_ for WM was 1.8–2.0 times larger for 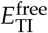 and 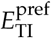 compared to tACS.

The optimal current patterns for *E*_ROI_ > 0.25 V/m (Fig. S11) were similar to the ones for maximal *E*_ROI_ (Fig. S10) except for 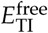 and 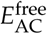 in the pallidum target: the two electrode pairs for *E*_ROI_ > 0.25 V/m are farther apart, which reduces both *E*_ROI_ and *Vol*_0.25_.

Figure 11 shows the optimal tTIS current patterns for *E*_ROI_ > 0.25 V/m in right pallidum (panel a) and the resulting field distributions (panel b). The 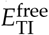 field (Fig. 11b1) shows a large area of high field strength in the right hemisphere’s WM. The stimulation produced a field with its peak area near the ROI, but the highest values were not at the target. This is consistent with the design of the study: the method we used minimized the stimulated volume, but it did not apply constraints that would restrict the field strength outside the ROI to be lower than that inside. While the optimal current pattern produced a field in the pallidum ROI of sufficient strength to achieve modulation, similar or stronger effects could result in other areas as well. The field for 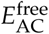 with the same *E*_ROI_ shows a small superficial region that receives lower field strengths than with tTIS, but the majority of the brain receives higher field strengths, with peaks in the WM inferior to the ROI (Fig. 11c1). The peak of the 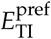 field was located closer to the ROI, with other small highfield areas nearby (Fig. 11b2), without affecting much of the superficial tissue between the ROI and the electrodes. The 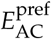 field looked similar, with slightly higher stimulation primarily in superficial areas superior to the ROI (Fig. 11c2).

**Figure 11:**
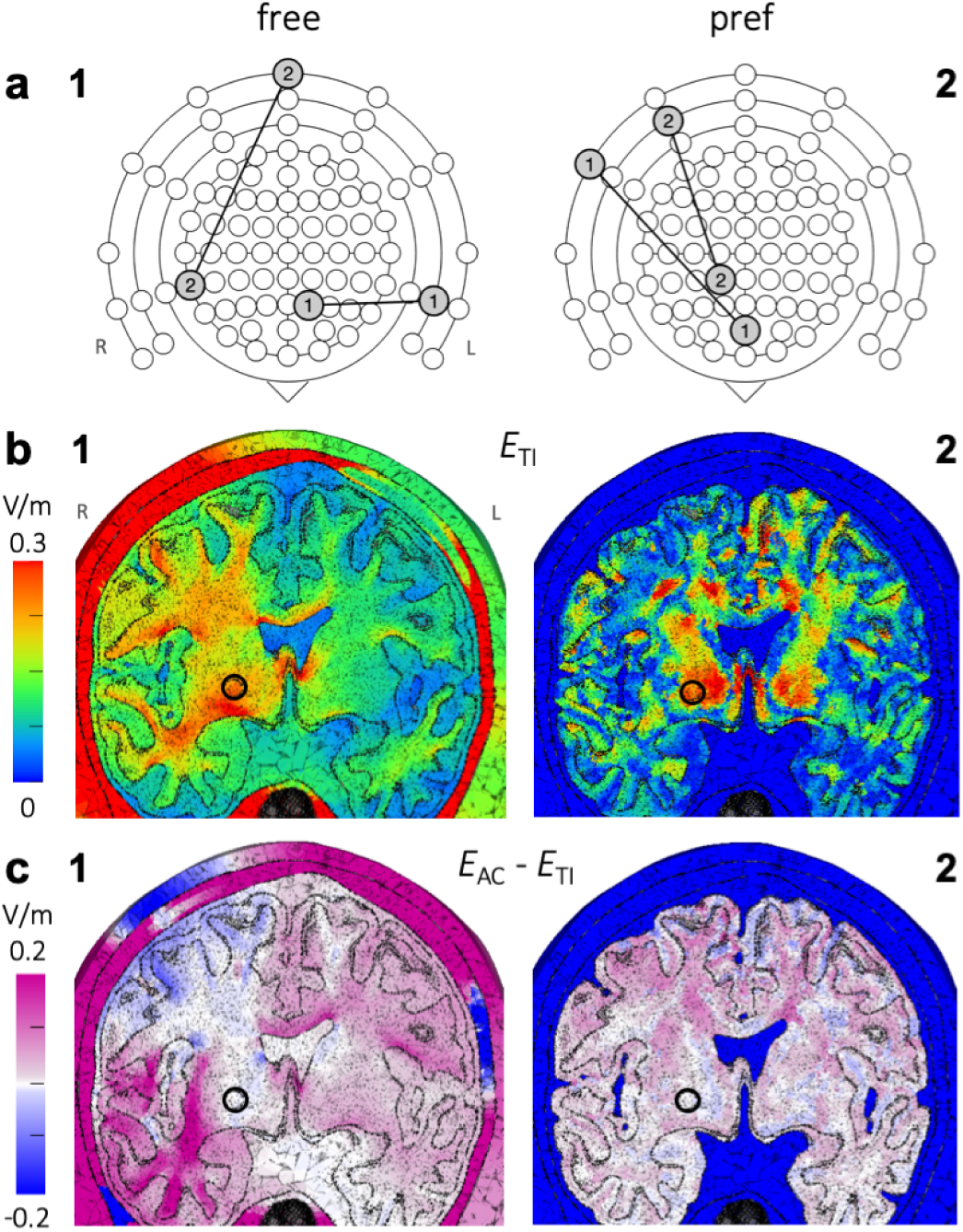
Study 3 – Optimal current patterns and electric fields for four-electrode tTIS and tACS in the right pallidum. **a)** Current patterns that minimized the stimulated brain volume (*Vol* _0.25_) while reaching a field strength of at least 0.25 V/m in a spherical ROI in the right pallidum for tTIS in a free (1) or preferred direction (2). Each electrode pair was represented by a line connecting two circles on an extended schematic of the 10-10 system; three rings were added around the standard schematic (Fig. 7a) to represent electrodes on the neck and cheeks (Fig. 10a) and the schematic was rotated 180 degrees to aid interpretation of panel b. For both current patterns, the input currents of each pair were equal (*I*_2_ = *I*_1_, *R* = 1). **b)** Optimal *E*_TI_ (fields resulting from the current patterns in panel a), and **c)** difference between optimal *E*_AC_ and optimal *E*_TI_, on a plane through the target region (indicated with a circle) for free (1) and preferred (2) directions, viewing towards the posterior direction (L and R in panel b1 indicate the left and right side of the head for all panels). Optimal current patterns for *E*_AC_ are shown in Fig. S11.

## 4 Discussion and Conclusions

### 4.1 Simulation of mice experiments

Our simulations with a murine head model showed that the stimulation parameters used in Grossman et al. (2017) produced simulated electric fields that were high enough to elicit neural firing. The peak area of the field was in the cortex and was lower and much smaller in spatial extent than that caused by tACS. Changing the ratio of input currents moved the peak across the cortex. Our results are consistent with the Grossman et al. experimental finding that tTIS can be used to selectively stimulate target regions in the mouse cortex at a suprathreshold level that is necessary to produce visible muscle twitching in the forepaws. We further note that our simulations showed that suprathreshold stimulation in mice should be achievable with tACS as well, which is in agreement with patch clamp experiments performed by Grossman et al., but the stimulation was much less focal and therefore targeted activation of specific muscles with tACS does not seem practical with a single configuration.

Our murine model did not have a skin layer and electrodes were placed directly on the skull, while in the Grossman et al. experiments electrodes were placed on shaved skin. Therefore, our simulations likely overestimated the field strengths. Given that the peak field is located near the top electrodes and that the skin of the mouse in this region is very thin, we do not expect this difference to strongly affect the results. Furthermore, as the maximum field strength (323 V/m) was far above threshold, overestimation due to the omission of the skin would not change our conclusion that muscle twitching can occur with these stimulation parameters. Grossman et al. included simulations with a murine FE model in their study, but the electrode parameters used did not match their experiments, and included tissue types were not specified (except for the exclusion of CSF, which was included in our model), making it difficult to compare their results to ours. Grossman et al. placed two cranial electrodes on their model at inter-electrode distances of 1.5 mm and 4.5 mm and two on the torso, with currents of 125 µA per pair, while in contrast 2.5 mm separation and 388 µA were used in their experiments and in our simulations. Scaling the results for their 1.5-mm simulations to 388 µA input current results in approximate peak values of 118 V/m for tTIS and 186 V/m for tACS, which is considerably lower than our results. This could be due to the larger distance between the two top electrodes in Grossman et al.’s simulations and the larger distance of the bottom electrodes to the top electrodes, likely combined with differences in the model, most notably the exclusion of CSF.

Both the spatial distribution and the values of the tTIS fields in the human model, as we discuss below, differed greatly from the results seen in the murine model. To achieve the same field strength in human motor cortex as was reached in the murine model, our simulations suggest that currents over 500 mA per electrode pair would be required. Thus while the mice experiments and simulations were informative, our results suggest that they are an inadequate model for predicting tTIS effects in humans due to the large difference in head morphology.

### 4.2 Implications of our study on the potential of tTIS in humans

Simulations of tTIS on a detailed human head model were performed with various configurations of four electrodes. Optimized field strengths for tTIS were comparable to those for conventional tACS in deep as well as superficial target areas. Median field strengths in any direction in ROIs in the hippocampus, pallidum and motor cortex were only slightly higher with tACS (0.25-0.60 V/m) than with tTIS (0.24-0.56 V/m) and both modalities produced simulated field strengths high enough to potentially achieve modulation in all three ROIs, in agreement with in vivo electric field measurements of tACS in humans (Huang et al., 2017; Chhatbar et al., 2018). These results suggest that deep areas of the brain could be stimulated with effects similar to those commonly found for conventional tACS in superficial areas, and that both tACS and tTIS could be used to achieve such modulation in deep as well as superficial areas.

However, our results also suggest that, in contrast to tACS, tTIS can produce electric fields that reach peak strength in deep brain areas. In particular this was achieved by placing two electrode pairs on opposite sides of the head. The peak could be steered towards superficial areas by changing the ratio of input currents, but this also decreased the peak’s field strength. Notably, the peak moved towards the electrode pair with the lower input current. Maximal target field strengths were achieved by placing the two electrode pairs next to each other, but this came at the expense of focality especially for deep targets: such configurations stimulated large areas outside of the ROI. These initial results suggest that tTIS field strengths can be maximized by placing electrode pairs next to each other, and focality can be maximized by placing pairs on opposite sides of the head, but more simulations will be needed to confirm this. Our initial optimization results for three ROIs showed that current patterns for tTIS can be found that are both theoretically effective (ROI field strength > 0.25 V/m) and focal (≤2% total brain stimulation).

Stimulation in the preferred direction (here defined as parallel to the dominant orientation of the neurons) also seems possible in both deep and superficial areas with tTIS as well as tACS; optimized field strengths in the pallidum and motor cortex ROIs were 0.26-0.31 V/m for tTIS and tACS. While the median ROI value of 0.17 V/m for tTIS in hippocampus did not reach the threshold of 0.25 V/m set in this paper, it should be noted that this was a conservative threshold, and the *maximum* value of 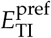 in the ROI was 0.36 V/m, suggesting that tTIS modulation in the preferred direction is likely to be possible in hippocampus as well. For the pallidum ROI, stimulation outside the ROI was generally smaller for the preferred direction as compared to the free-direction case, and the highest field strengths were reached closer to the ROI, suggesting the opportunity for greater focality if indeed modulatory effects are dominated by the field component in the preferred direction. However, for hippocampus and M1, stimulation focality was poorer in the preferred direction. Optimized simulations for a much richer set of targets in the future will be needed to understand these differences.

While similar field strengths could be reached with tTIS and tACS in deep as well as superficial targets, there are several notable differences between the two modalities. First, as described above, tTIS can achieve peaks in deep areas while not stimulating overlying areas, which is not possible with tACS. Second, the volume of brain tissue that was stimulated when targeting a deep region was much larger with tACS, for standard as well as optimized current patterns. Although tACS fields can be made more focal by optimizing configurations with more than four electrodes, in principle the same would be true for tTIS. Third, fields for tTIS were affected much more by the current ratio, allowing for more control of the field distribution via the same electrode montage. This could prove highly useful in practice when one wants to reach multiple brain areas in one session without adjusting electrodes, or one wants to be able to adjust the focus of stimulation online based on some measurement. Finally, for configurations with pairs on opposite sides of the head, effects of current ratio on the maximum and stimulated volume were opposite for tTIS and tACS, and the directions of maximum electric fields were perpendicular in certain areas of the brain. This suggests there might be circumstances under which either tTIS or tACS is better suited to achieve the desired goal and thus having both options broadens the utility of tCS.

We evaluated 146M input current patterns with four electrodes and the maximum tTIS field strength reached anywhere in the brain and in any direction was 0.77 V/m. With this comprehensive set of configurations, it is unlikely that much higher field strengths can be achieved with tTIS in the human head with a total current of 2 mA. Even with sophisticated optimization methods, the tTIS field strength will always be bounded by the summed field strengths of the input fields, which, with a total injected current of 2 mA, will not reach close to the ∼28– 120 V/m^6^ generally thought to be necessary to cause supratheshold excitation in human corticomotor neurons (Radman et al., 2009b). An input current of 38 mA per electrode pair would be needed to reach 28 V/m in the motor cortex of our human model. We thus conclude that it is likely not possible to achieve suprathreshold stimulation in humans with tTIS at safe input currents. We note that simulations of DBS show field strengths around 75 V/m (Åström et al., 2015). Therefore, tTIS does not seem to be viable as a non-invasive alternative to DBS in humans.

In three of our studies (2a,2b,2c), we systematically changed electrode placement and observed the effects on electric field distributions. We were able to draw some general conclusions from these studies; however the results were non-trivially related to anatomical details in the model. Thus we intend to carry out further studies using several head models and a larger set of anatomical targets. Our goal is to determine if there are generalizable recommendations across models and targets, or if instead detailed optimization is needed for each intended application.

### 4.3 Comparison to other recent studies

Grossman et al. performed FE simulations with a cylindrical and a spherical head model for several current ratios, which they verified with measurements in a phantom. As a control of our implementation, we repeated their sphere simulations and results (not shown) looked similar. In addition, as noted above, our murine simulation of their experimental results in mice was consistent with their report, which substantiates the veracity of our simulation methods.

Huang and Parra (2019) reported the first simulations of tTIS with a realistic human head model using two configurations with *R* = 1 and *R* = 4. They found that for these configurations with two pairs on opposite sides of the head in an axial plane, the tTIS peak occurred deep in the brain. For comparison purposes, we performed additional simulations with the configurations they used and assumed some parameters not described in their report (results in Fig. S12; compare to Fig. 2 in Huang and Parra (2019)). The two sets of simulations show globally similar results with peaks in the same areas, mostly near the lateral ventricles for tTIS and superficial for tACS, but local distributions are different and field strengths are higher in our results. This may be due in particular to differences in head geometry and conductivities; especially for tACS large differences in posterior areas can be seen that are likely due to inclusion of anisotropy in our model. The authors of Huang and Parra (2019) found that field strengths for tTIS were lower than for tACS throughout the brain, which is in agreement with our results, and therefore concluded that tTIS does not have benefits compared to tACS. In contrast, our results suggest that tTIS might provide benefits over tACS in the form of increased focality and steerability. Furthermore, when optimization was used, the target field strengths for tTIS were almost identical to those for tACS.

Another approach with potential for focused deep stimulation from transcranial electrodes was reported in Vöröslakos et al. (2018). In this study, interleaved short pulses from different electrodes were used to achieve deep stimulation, relying on membrane time constants that are longer than the pulse duration to effectively superimpose the temporally distinct pulses at depth. As pointed out in Huang and Parra (2019), from a modeling point of view, as long as neuronal response models are not considered, this is equivalent to modeling any other direct transcranial simulation including tDCS and tACS.

### 4.4 Effects of head model parameters on simulations

We did extensive comparisons, not reported here, to elucidate how tTIS simulations are effected by modeling parameters such as the inclusion of anisotropy, details of the lateral ventricles, and skull conductivity inhomogeneity. In general we found that both white matter anisotropy and accurate modeling of the ventricles had substantial impact on the field distributions in our results. For maximum feasible verisimilitude, in this report we used a model that included both of those properties. Since we mostly evaluated tTIS fields deep in the brain, individual differences in cortical folding likely matter less here than for conventional tCS simulations. A detailed study of the relative importance of model parameters and inter-subject variability might be informative and useful to optimize effort in future tTIS modeling studies but is outside the scope of this initial report, which focuses on using an extensive set of stimulation parameters on a single model to better understand the limits and parameters of tTIS fields compared to those from tACS.

### 4.5 tTIS field strength and neuromodulatory effect

Our simulations give a thorough insight into the electric field distributions during tTIS, and specifically how and where temporal interference effects occur. However, we do not study how these fields would affect neurons, neuronal populations, or neural circuits. Practically achievable temporal interference fields at the difference frequency were strong enough to achieve spiking activity in mice in Grossman et al. (2017), and would theoretically drive subthreshold neuromodulation in humans. However, more research is needed into the mechanisms by which mouse neurons spike in response to the difference frequency given the omnipresent carrier frequency, and whether those same mechanisms can drive subthreshold membrane responses in human neurons. The Huang and Parra (2019) paper, like ours, implicitly assumes that modulation depth is the driving mechanism but we believe that is yet to be determined. Neuron models could provide insight into the prospects and mechanisms of tTIS on a neural level, similar to what has been done for tDCS (Reato et al., 2010; Rahman et al., 2013). Initial results for tTIS using a spherical volume conductor and Hodgin-Huxley point neuron model recently reported in an unpublished manuscript posted on bioRxiv (Cao and Grover, 2017) suggest a complex dependence on carrier frequency, difference frequency, and the number of mutually interfering fields. We are currently working on combining our realistic FE simulation results with a Hodgin-Huxley style cable model to study this central question of the tTIS mechanism of neuromodulation.

### 4.6 Optimization of tTIS

Using an exhaustive search over nearly 146M current patterns (7M electrode configurations with 21 current ratios for each), we found Pareto-optimal four-electrode current patterns for maximal stimulation in three ROIs with minimal stimulation of regions outside the ROI. Any current pattern on these boundaries is optimal for the definition used here. Dependent on the requirements of a planned experiment, one would select a point on these lines that achieves the desired target field strength and focality. For 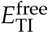, maximal ROI field strengths were 2.9– 5.4 times larger than with the configuration based on Grossman et al. (2017); for equal field strengths the stimulated volume outside the ROI was smaller. Although this method was successful at maximizing field strength in a specific region while minimizing the total brain volume stimulated above certain limits, large areas outside of the ROI were still stimulated. This could potentially be reduced by extending our approach with additional constraints such as a limit on the maximum field strength outside the ROI. Methods to further shape the field could include applying tTIS with more than two electrodes per frequency and/or more than two stimulation frequencies (Cao and Grover, 2017). In this case, the exhaustive search approach would quickly become unfeasible.

Mathematical optimization algorithms could be devised to solve such problems in a reasonable time frame, as has been done for example for tDCS (Dmochowski et al., 2011; Ruffini et al., 2014; Guler et al., 2016). These kinds of algorithms also allow for more sophisticated constraints to minimize fields in specific areas and to optimize for multifocal targets (e.g., based on fMRI). However, compared to tDCS (or tACS with a quasi-static model as used here and elsewhere), where the stimulation effect depends linearly on the injected current, tTIS presents a much more difficult set of challenges, as the effect at any location is driven by the locally weakest field and the alternating nature of the current makes the stimulation invariant to the polarity of the input currents (distinct from tDCS). This results in a non-linear and nonconvex optimization problem that substantially increases the challenge of optimizing the electrode currents to maximize the tTIS effect. Our team is currently working on a method to overcome this challenge by developing a set of sufficient conditions derived from the mathematical structure of the problem, that are both reasonable and can be checked in practice, that will allow us to guarantee the global optimality of a locally optimal convex relaxation. The results described here are critical to inform sensible constraints and structure to any future attempts at mathematical optimization.

### 4.7 Additional limitations of our results to date

In addition to the considerations of modeling parameters, inter-subject variability, neuromodulatory effect, and optimization methods discussed above, we also acknowledge that we only considered three potential targets among a large number of regions of potential clinical interest. Furthermore, we considered those regions as isolated targets, not in combinations or as part of a network. We believe that these are all topics of interest for future research, both from a modeling point of view, and more importantly, in collaboration with experimental investigations of tTIS effects. Only through a series of such studies can we answer the many open questions regarding the potential utility of this modulation approach in practice.

### 4.8 Conclusions

According to our biophysical models, transcranial temporal interference stimulation can be used for suprathreshold stimulation in small animals, but not in humans. Field strengths achieved in humans were in the range of those previously reported with conventional tACS. Major differences between tTIS and tACS are that with tTIS 1) for targets deep in the brain, overlying areas are stimulated less, and 2) the peak of the field can be steered to a desired location using a single electrode montage by varying the input current at each frequency. We conclude that tTIS has the potential to be a more focal and steerable alternative to tACS for neuromodulation of deep brain areas. To better understand the working mechanisms and prospects of tTIS in humans, more research is needed utilizing combinations of FE simulations, neuron models, and experiments. Our future work will be focused on mathematical optimization of tTIS, neuron models, and simulations investigating additional brain areas of clinical or scientific interest with multiple head models.

## Acknowledgements

This project, and in particular SR and DHB were supported in part by the National Institute of General Medical Sciences of the National Institutes of Health under grant number P41 GM103545-18. ADD was supported in part by the National Science Foundation under CAREER 1351112. The authors would like to thank Magdalena Schwarzl for providing the code that produced Figure 6c.

## Footnotes

1 also called “interferential stimulation”, “interference currents” or “interferential current therapy”

2 Although tissue conductivity is frequency dependent and tTIS involves frequencies that are much higher than those used for tACS, recent tTIS simulations reported qualitatively similar results for standard conductivity values at 0 versus 1 kHz (Huang and Parra, 2019), so we used conductivity values commonly used for tCS simulations.

3 In Grossman et al. (2017), *I*_L_ and *I*_R_ were labeled *I*_2_ and *I*_1_, respectively.

4 Note that the labels of the fields only refer to the side of the head on which the respective electrodes are placed; the field itself spreads throughout the entire head.

5 When describing the peak or maximum for tTIS, this always refers to the tTIS field strength calculated from Eq. 1; the two high-frequency fields have their maxima in other areas (see Fig. S1 for an example).

6 These values were determined using square pulse step currents and may differ for sinusoidal current.

## Notes

#### Summary of Updates

Study 3: 'Optimization of four-electrode tTIS' was updated to optimize current patterns for both target field strength and focality, as opposed to target field strength alone, as reported in the first version of the paper (changes in Sections 2.8, 2.9, 3.4 and Supplementary Material). Additional minor edits to text and figures for clarity.

