## Supplementary Material for "Prospects for transcranial temporal interference stimulation in humans: a computational study"

In this document, we present several figures and animations that complement the results described in the paper. Animated figures are used to display results for various current ratios or configurations in one figure to aid the evaluation of the effect of either parameter. For this reason, this document is best viewed in Adobe Reader, which will display the animations as intended. Any other viewer will show a still figure in place of each animation. In this case, the displayed figure presents the results for the median value of the parameter, i.e., either $R = 1$ or the configuration with the displaced electrodes at the center of the range. The parameter space was subsampled in order to reduce the size of the document: for all figures that animate the current ratio, the values used for $R$ are 11 logarithmically spaced values between 0.1 and 10; for figures that animate the electrode configuration, the values used for Study 2b are 17 linearly spaced values between -80 and 80 degrees and the values for Study 2c are 8 linearly spaced values between 15 and 155 cm. Controls under each image allow for pausing, stepping through the animation frame by frame, and adjusting the frame rate. If the animation does not start immediately, it may help to scroll to the next page and back.

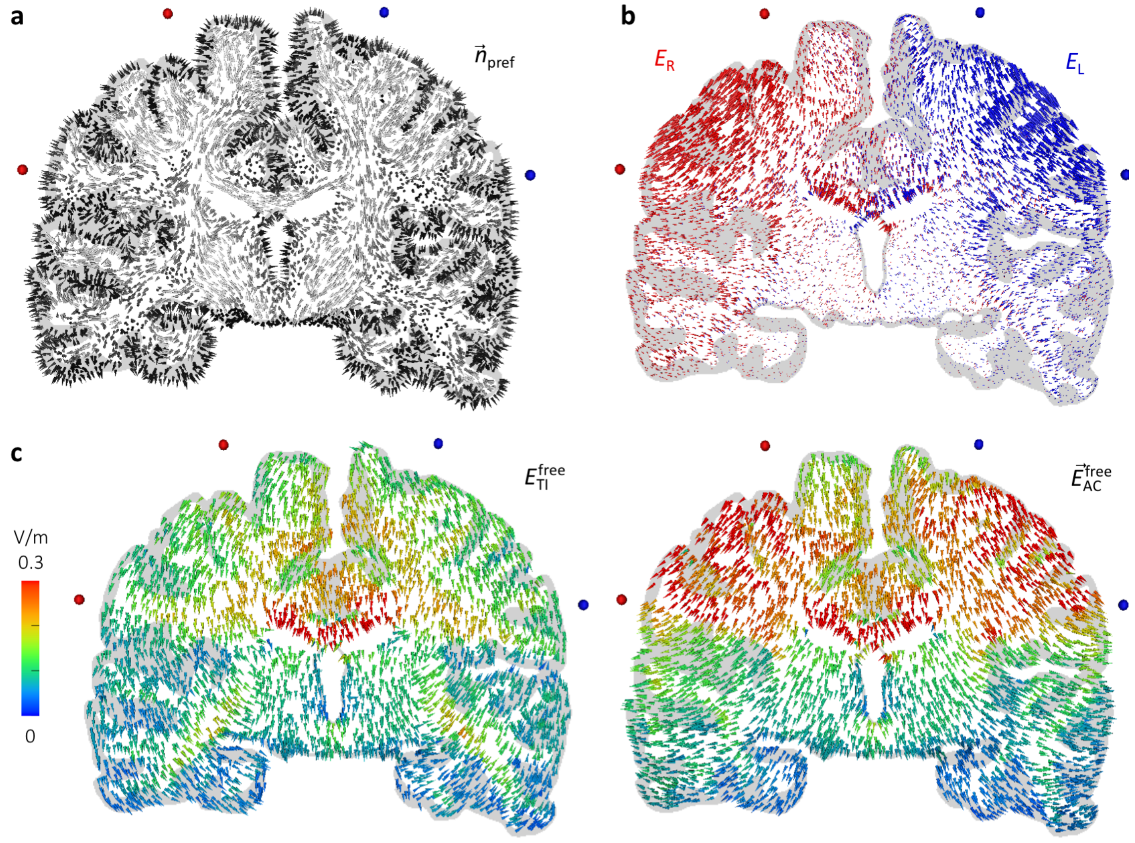


**Figure S1: Study 1 – Directions of preferred and simulated electric fields.** Displayed are electric field vectors on a plane through the electrodes (all in the coronal plane), viewing towards the posterior direction (left side of brain on right side of image). Red and blue spheres indicate the locations of the electrodes that provide $I_R$ and $I_L$, respectively. **a)** The preferred direction vector field $\vec{n}_{\text{pref}}$ was constructed from DTI data in white matter (light gray cones) and vectors perpendicular to the gray and white matter boundaries in gray matter (dark gray cones). This field was used to calculate $E_{\text{TI}}^{\text{pref}}$ (Eq. 3) and $E_{\text{AC}}^{\text{pref}}$ (Eq. 4) for all analyses. **b)** The two fields produced by currents $I_L = I_R$, with field strength and direction represented by cone size and direction, respectively. **c)** Simulated electric fields for TIS and tACS with $I_L = I_R = 1$ mA. For TIS, the direction of each cone is the direction that maximizes the field strength in that element, i.e. the $\vec{n}$ that maximizes $\vec{E}_{\text{TI}}$ in Eq. 1; the color of the cone represents the field strength in that direction ($E_{\text{TI}}^{\text{free}}$). For tACS, the cone direction and color represent the direction and field strength of $\vec{E}_{\text{AC}}$ (Eq. 4).

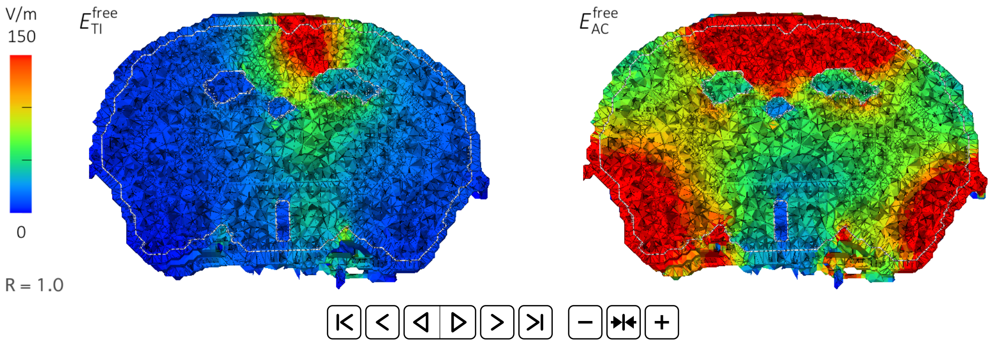


**Figure S2: Field strengths for TIS and tACS in the murine model for various current ratios.** Displayed is electric field strength on a plane through the electrodes, looking towards the anterior direction (left side of brain on left side of image), for $R = I_R/I_L$ from 0.1 to 10 (animated) with $I_R + I_L = 0.776$ mA. The distinct areas with low field strengths are the fluid-filled ventricles. Electrode placement was based on experiments by Grossman et al. (2017).

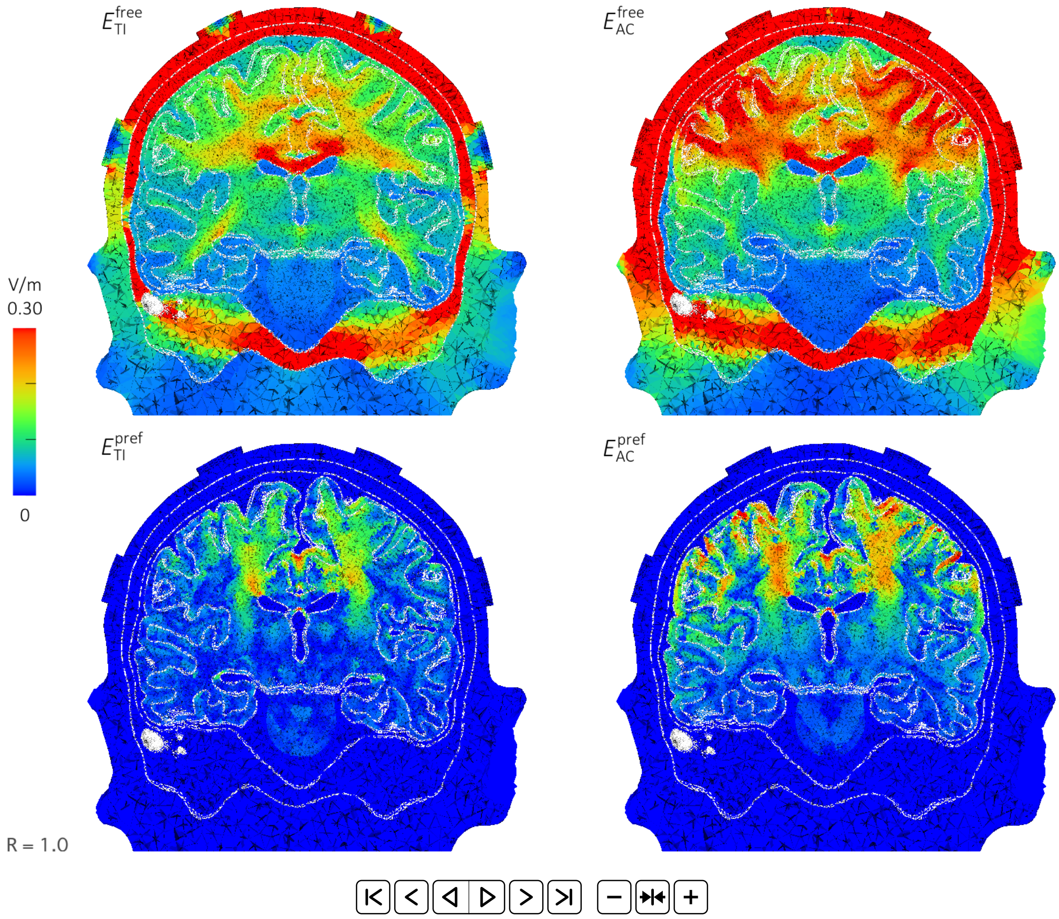


**Figure S3: Study 1 – Field strength distributions for TIS and tACS with various current ratios.** Displayed are field strengths on a plane through the electrodes (all in the coronal plane), viewing towards the posterior direction (left side of brain on right side of image), for $R = I_R/I_L$ from 0.1 to 10 (animated) with $I_R + I_L = 2$ mA. The preferred direction is only defined for brain elements; hence for plots of $E_{TI}^{pref}$ and $E_{AC}^{pref}$, all non-brain elements have a value of 0. A subset of these figures was shown in Fig. 5.

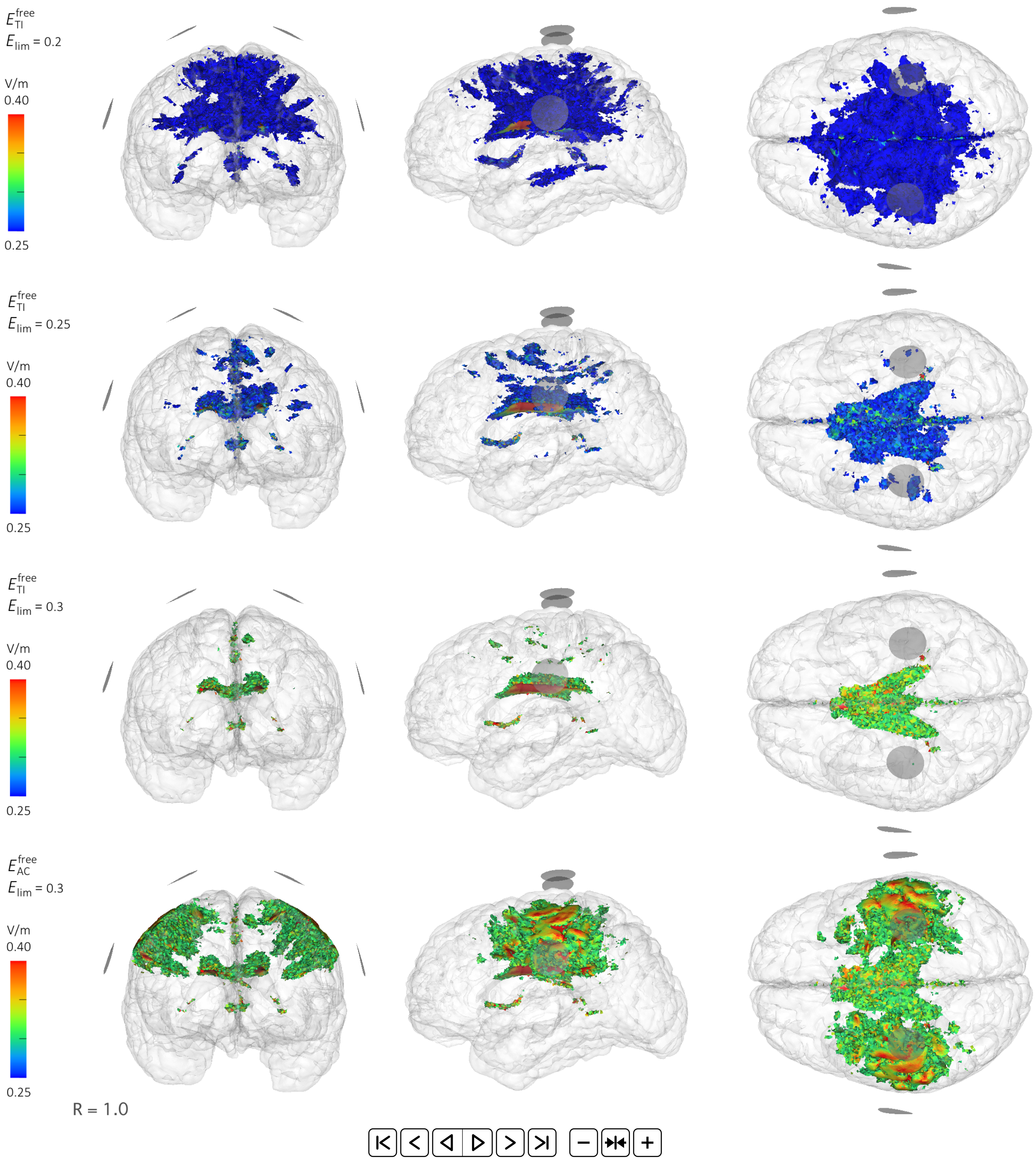


**Figure S4: Study 1 – Stimulated brain volume for TIS and tACS with various current ratios and $E_{\text{lim}}$ values.** Each figure shows the brain volume for which $E > E_{\text{lim}}$ for current ratios $R = I_R/I_L$ from 0.1 to 10 (animated) with $I_R + I_L = 2$ mA. Rows 1-3: $E_{\text{TI}}^{\text{free}}$ with $E_{\text{lim}}$ = 0.2, 0.25 and 0.3 V/m, respectively. Row 4: $E_{\text{AC}}^{\text{free}}$ with $E_{\text{lim}}$ = 0.3 V/m. From left to right, the brain is visualized from the front, left and top; all images are on the same scale. The configuration consists of four electrodes in the coronal plane; electrode surfaces are visualized as gray disks. A subset of these figures was shown in Fig. 6a.

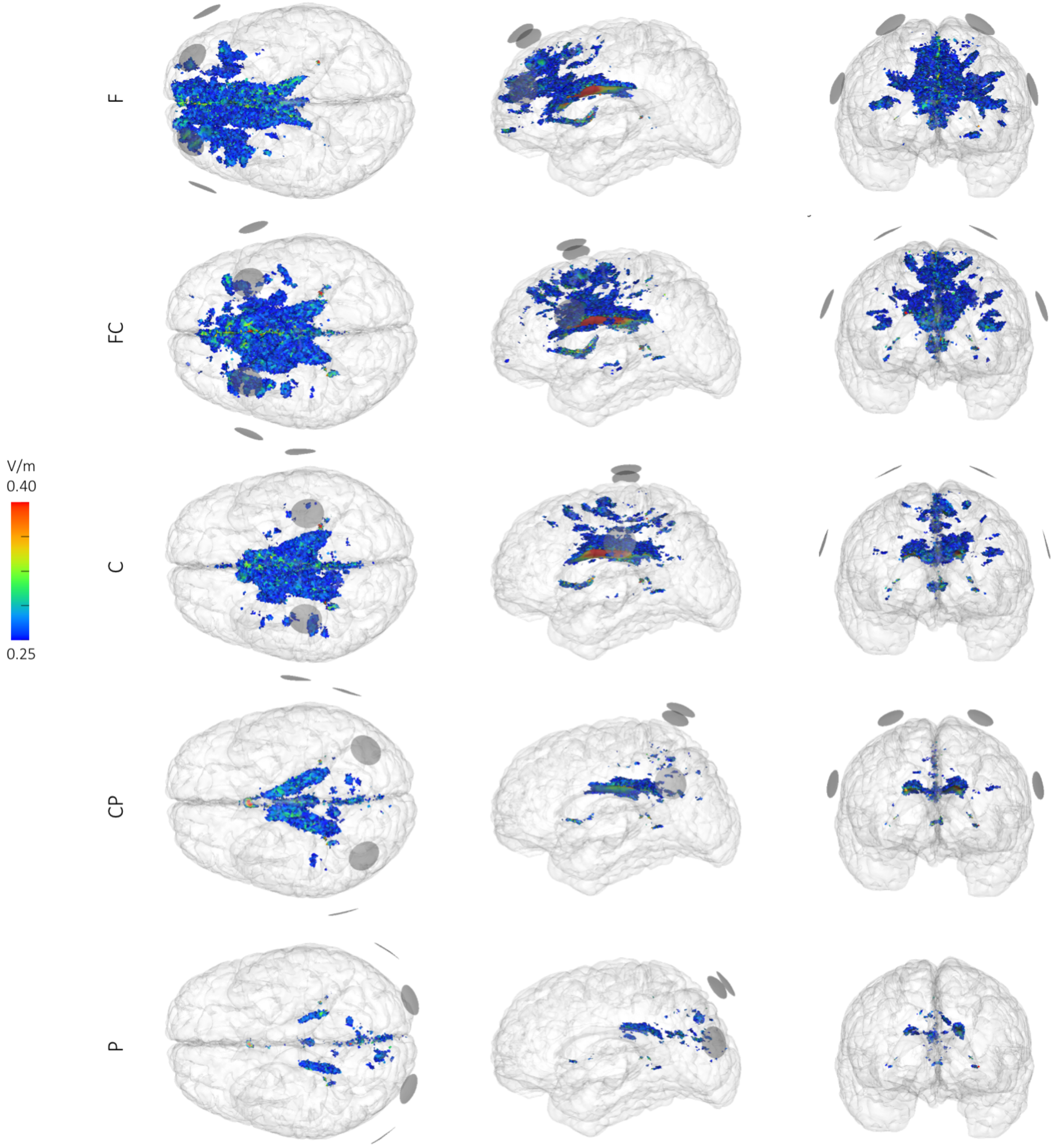


**Figure S5: Study 2a – Stimulated brain volume for $E_{TI}^{free} > 0.25$ V/m for five standardized electrode configurations with $I_R = I_L = 1$ mA.** From left to right, the brain is visualized from the front, left and top; all images are on the same scale. Displayed from top to bottom are configurations with electrodes through the F1, FC1, C1, CP1 or P1 locations of the 10-10 electrode system; electrode surfaces are visualized as gray disks.

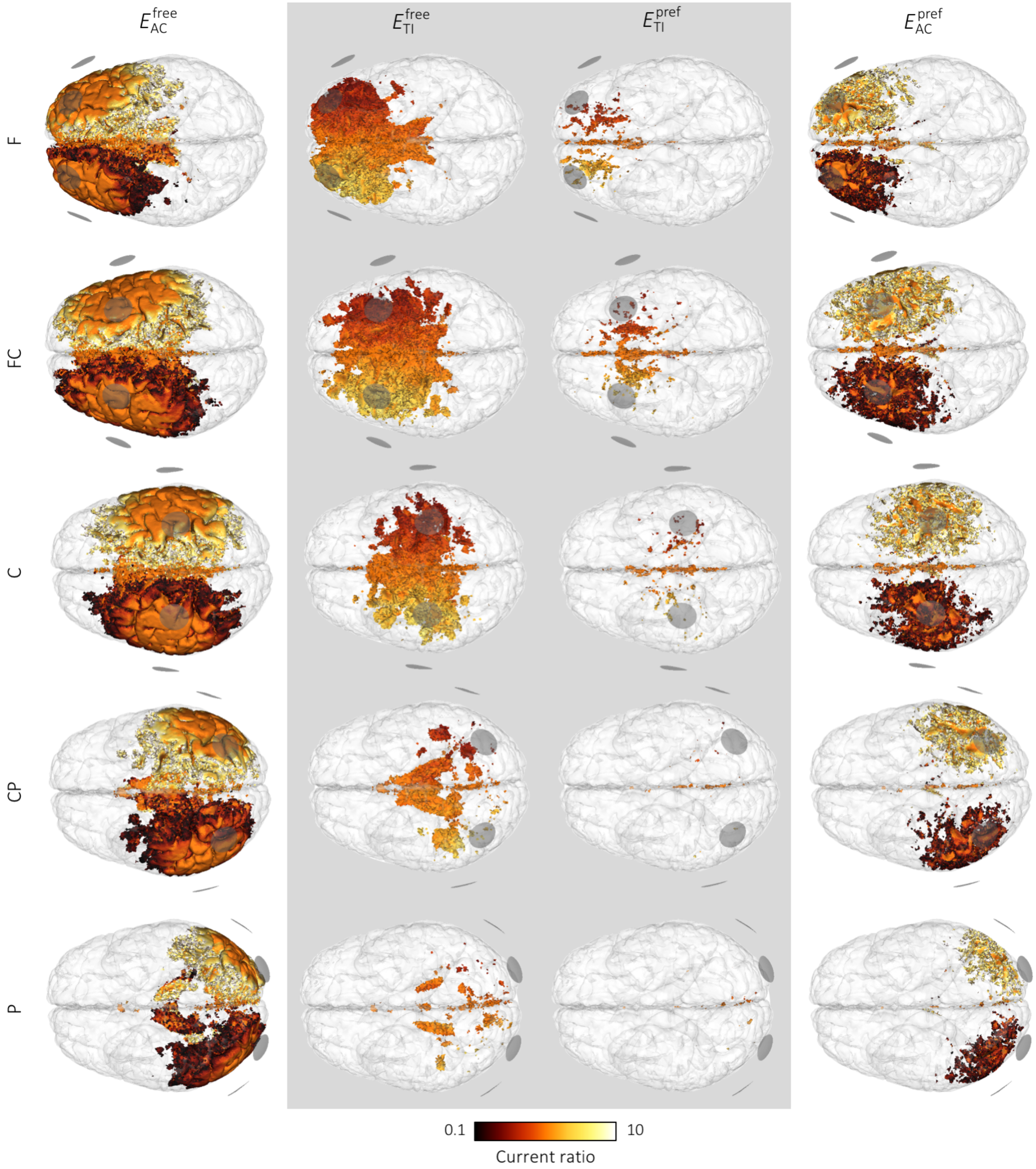


**Figure S6: Study 2a – Stimulated brain volume for TIS and tACS with five standardized electrode configurations with various current ratios.** Colors represent the current ratio $R$ that stimulates the brain above 0.25 V/m at that location; if this is true for multiple ratios at a given location, the ratio closest to 1 was selected for display. The combined volume in one image thus represents the total brain volume that can be stimulated with this configuration, i.e. all parts of the brain that can be reached by varying the input currents. Displayed from top to bottom are configurations with electrodes through the F1, FC1, C1, CP1 or P1 locations of the 10-10 electrode system. From left to right, $E_{TI}^{free}$, $E_{TI}^{pref}$, $E_{AC}^{free}$ and $E_{AC}^{pref}$ are shown.

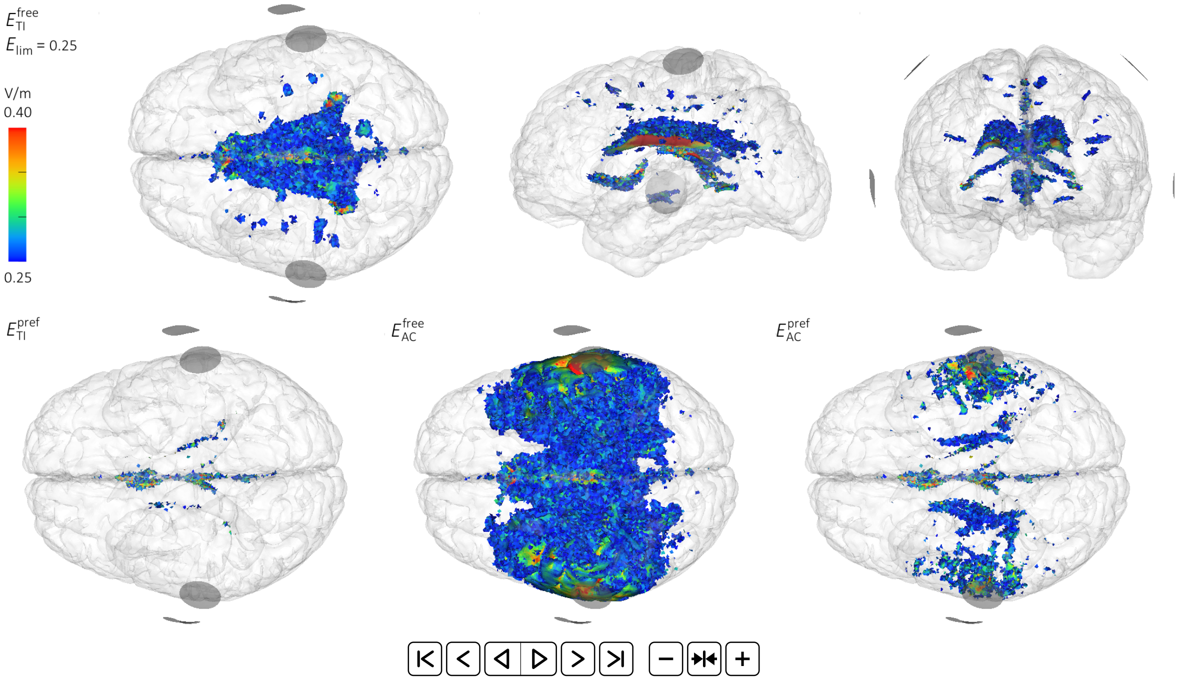


**Figure S7: Study 2b – Stimulated brain volume for TIS and tACS with a densely sampled set of parallel configurations.** Each figure shows the brain volume for which $E > 0.25$ V/m for a series of configurations with four electrodes that are moved from the front towards the back of the head (animated) and $I_R = I_L = 1$ mA; electrode surfaces are visualized as gray disks. Top row: top, left and front view for $E_{TI}^{free}$. Bottom row: top view for $E_{TI}^{pref}$, $E_{AC}^{free}$ and $E_{AC}^{pref}$.

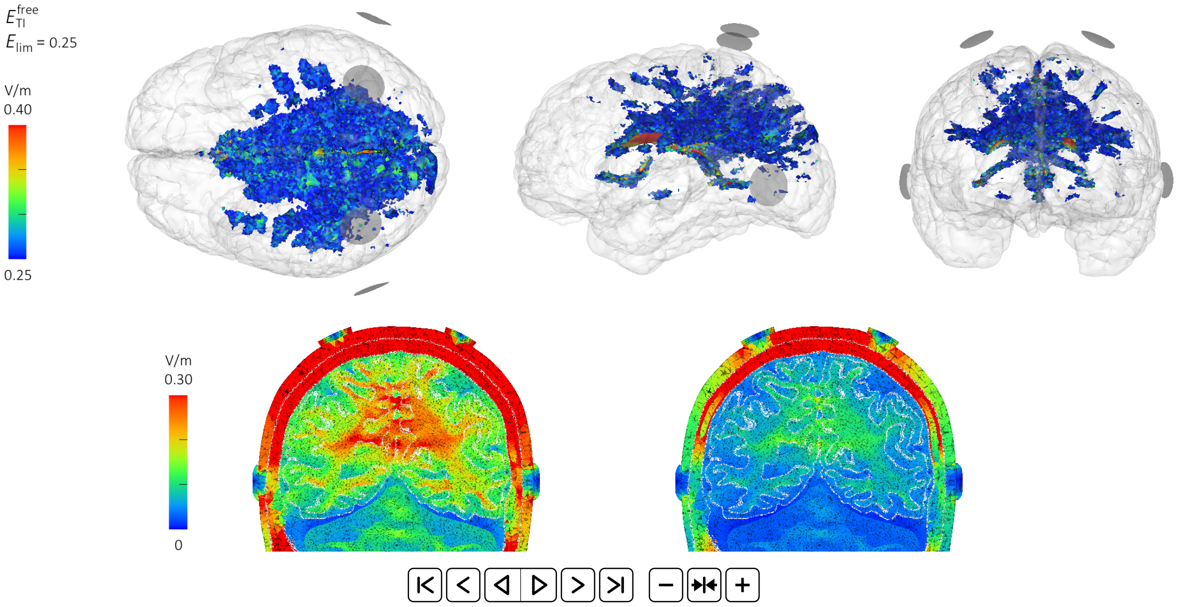


**Figure S8: Study 2c – Results for TIS with electrode positions varied vertically with $I_R = I_L = 1$ mA.** Top: Brain volume for which $E_{TI}^{free} > 0.25$ V/m. The brain is visualized from the top, left and front; all images are on the same scale; electrode surfaces are visualized as gray disks. Bottom: Field strengths on a plane through the electrodes (plane through CP1 and CP2), viewing towards the posterior direction. The left figure shows absolute field strength (similar to previous figures); for the right figure, field strengths for each configuration were normalized to the maximum field strength for that configuration.

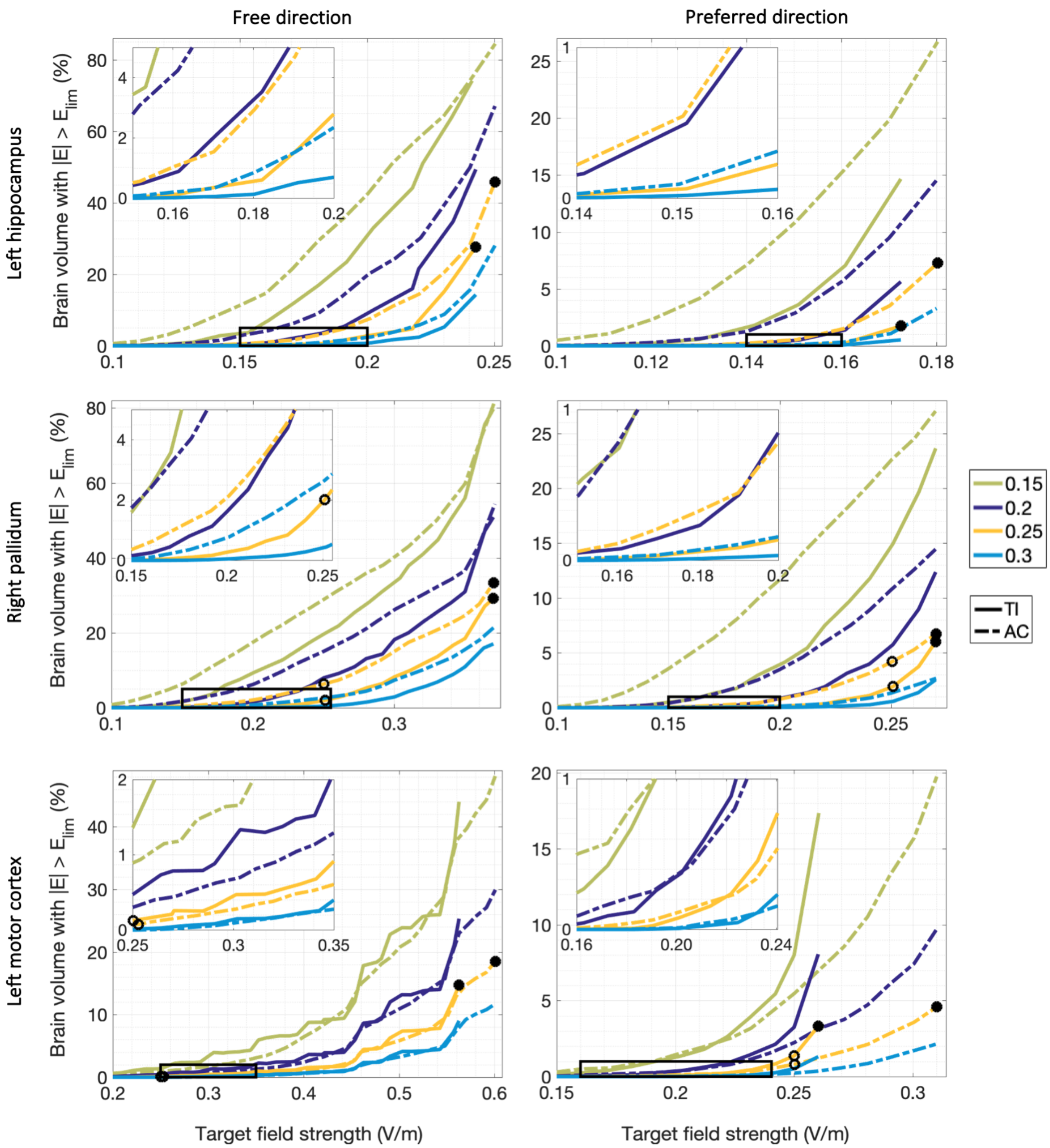


**Figure S9:** **Study 3 – Pareto boundaries of optimal field strength and focality achievable with four-electrode current patterns in three target areas.** From a set of 146M available current patterns, these plots present the lowest achievable stimulated brain volumes for four stimulation limits as a function of median field strength in a small region of interest (ROI). From these lines, the most suitable current pattern can be selected to achieve a specific experimental goal. Each row presents results for one ROI; the left and right columns present results for stimulation in any direction or along the dominant direction of the neurons, respectively. Red boxes indicate the locations of the insets. Black circles indicate current patterns that minimized the brain volume stimulated over 0.25 V/m while either maximizing field strength in the ROI (filled) or reaching at least 0.25 V/m in the ROI (open); these current patterns are visualized in Figs. S10 and S11, respectively.

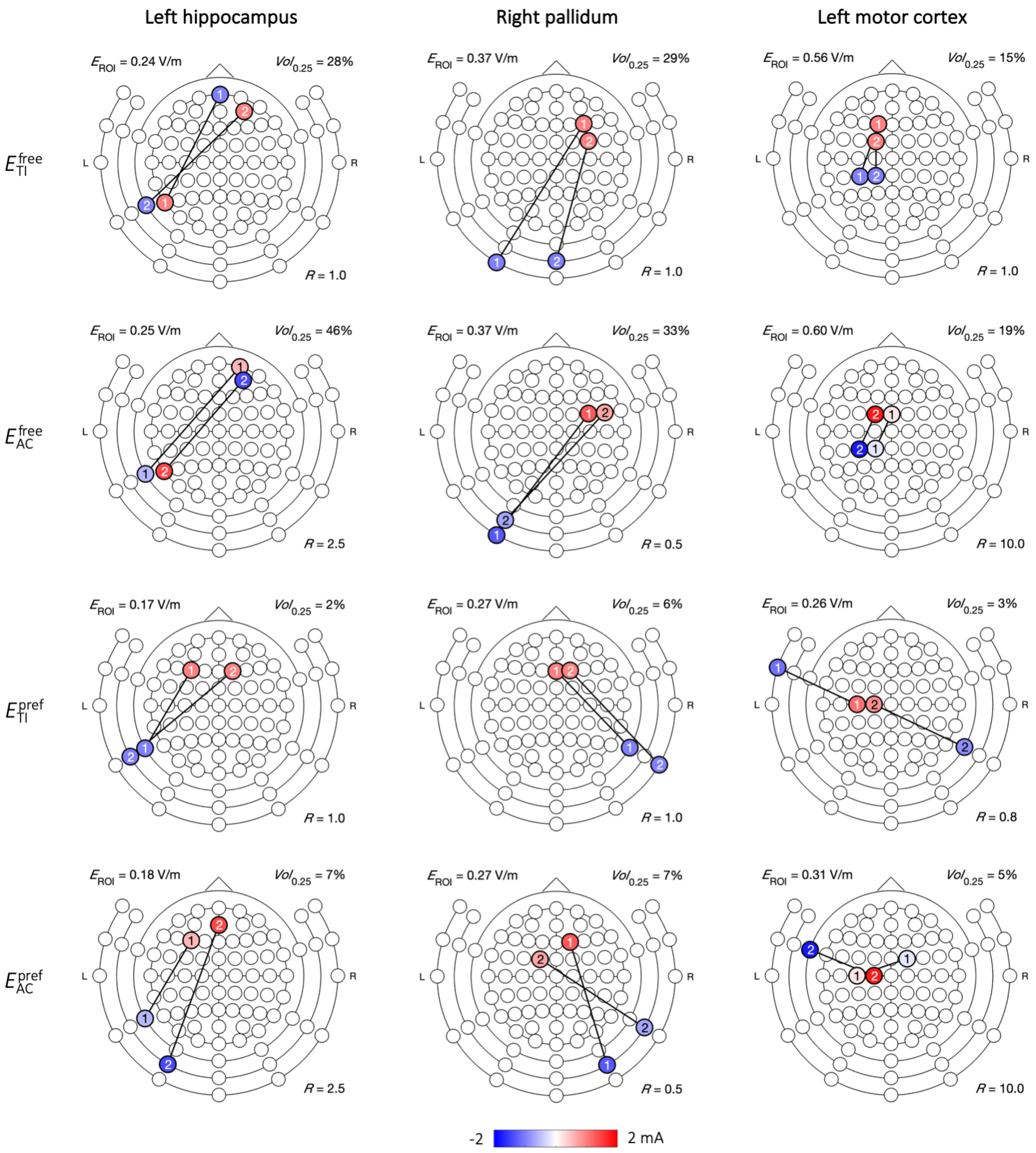


**Figure S10: Study 3 – Optimal four-electrode current patterns for maximal field strength in three target areas.** From a set of 146M current patterns (constructed from 7M configurations of 4 out of 88 electrode locations (Fig. 10a) and 21 current ratios), the current patterns were selected that minimized the brain volume stimulated over 0.25 V/m ($Vol_{0.25}$) while achieving maximal median field strength ($E_{ROI}$) in a small target area, i.e., the highest field strength on the Pareto boundary (Fig. S9, filled black circles). Results are displayed on an extended version of the standard 10-10 schematic (Fig. 7a), where the two electrode pairs (one for $I_1$, oscillating with $f_1$, and one for $I_2$ oscillating with $f_2$) of the best-performing configuration were colored with the current (mA) that produced the optimal result; the current ratio $R$ is indicated in each plot. The electrodes were colored as anodes (red) and cathodes (blue) for clarity, but for tTIS the selection is arbitrary and for tACS only the phase of the two pairs with respect to each other is relevant.

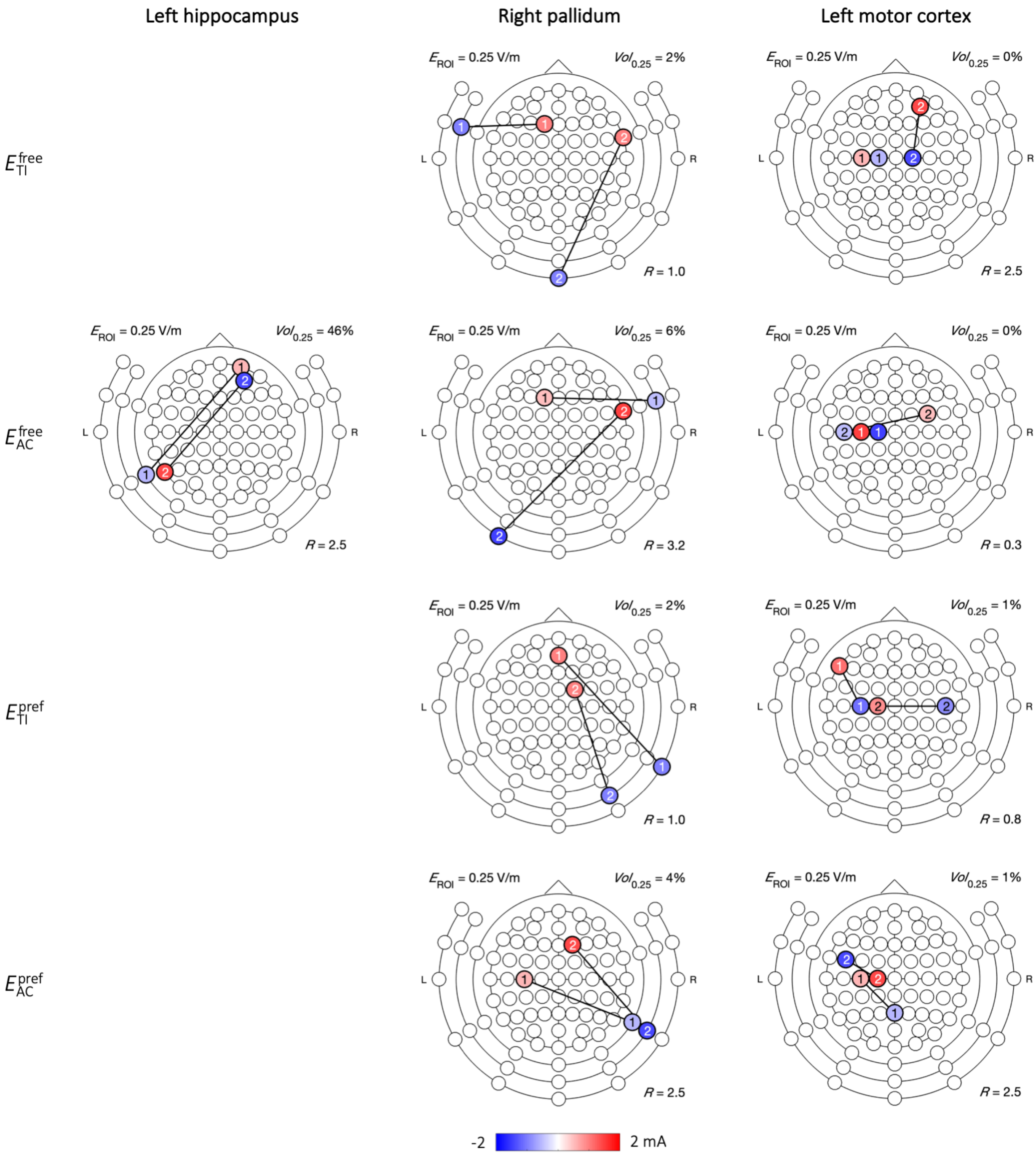


**Figure S11: Study 3 – Optimal four-electrode current patterns to reach at least 0.25 V/m in three target areas.** From a set of 146M current patterns (constructed from 7M configurations of 4 out of 88 electrode locations (Fig. 10a) and 21 current ratios), the current patterns were selected that minimized the brain volume stimulated over 0.25 V/m ($Vol_{0.25}$) while achieving a median field strength ($E_{\mathrm{ROI}}$) of at least 0.25 V/m in a small target area (Fig. S9, open black circles). For the hippocampus target, $E_{\mathrm{TI}}^{\mathrm{free}}$, $E_{\mathrm{TI}}^{\mathrm{pref}}$ and $E_{\mathrm{AC}}^{\mathrm{pref}}$ did not reach this limit and hence do not have a plot. See caption for Fig. S10 for further details on the visualization.

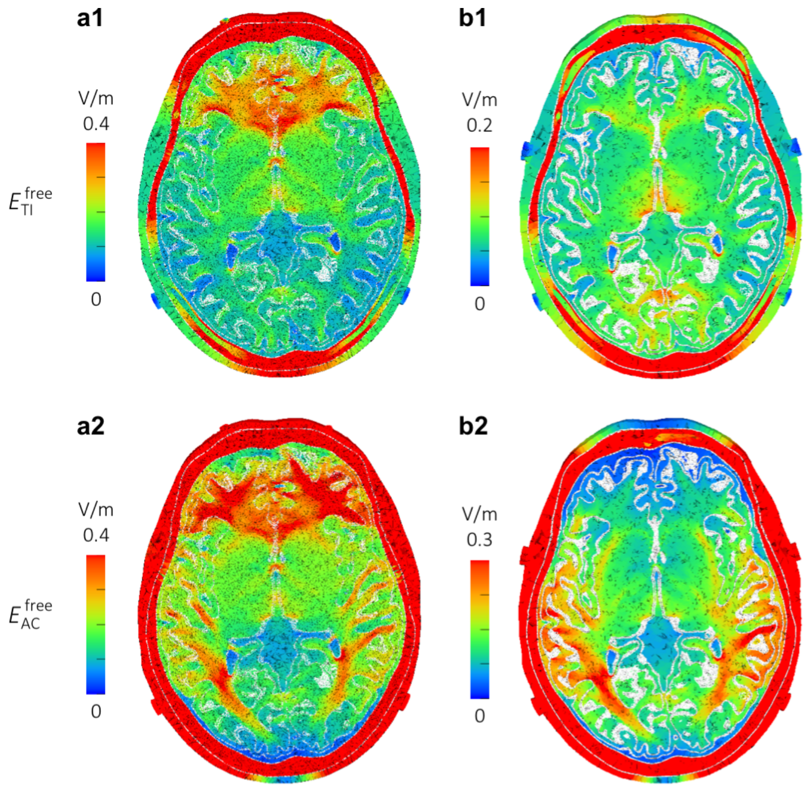


**Figure S12: Simulations based on Huang and Parra (2019).** Using the head model described in Section 2.2, we performed simulations with electrodes of 0.5 mm radius at **a)** FP1 and P7 for $I_L$ and FP2 and P8 for $I_R$, or **b)** FT7 and P7 for $I_L$ and FT8 and P8 for $I_R$, with $I_L = I_R = 1$ mA. Displayed are field strengths on a plane through the electrodes (all in an axial plane) viewing towards the inferior direction for 1) $E_{TI}^{free}$ and 2) $E_{AC}^{free}$. Panels a and b can be compared to the third column of Figs. 2B and 2C, respectively, in Huang and Parra (2019), where $E_{TI}^{free}$ and $E_{AC}^{free}$ are labeled "interferential" and "conventional" stimulation, respectively. Color bar limits were set to the values used in Huang and Parra (2019).
